# A Telltale Sign of Irreversibility in Transcriptional Regulation

**DOI:** 10.1101/2022.06.27.497819

**Authors:** Robert Shelansky, Sara Abrahamsson, Michael Doody, Christopher R. Brown, Heta P. Patel, Tineke L. Lenstra, Daniel R. Larson, Hinrich Boeger

## Abstract

Transcription occurs in stochastic bursts, i.e., transcription events are temporally clustered. The clustering does not ensue from environmental fluctuations but springs from the intrinsically stochastic behavior of the regulatory process that controls transcription. Based on microscopic observations of transcription at a single gene copy of yeast, we show that the regulatory process is cyclic and irreversible, i.e., the process violates the detailed balance conditions for thermodynamic equilibrium. The theoretical significance of this finding is discussed.

## Introduction

The nucleosome, which may form on any DNA sequence, albeit with varying propensity ^1^, serves as a general repressor of transcription initiation in the eukaryotic cell ^2^. Transcriptional activation is accompanied, therefore, by the removal of nucleosomes from promoter sequences ^3^. However, the removal remains incomplete at any position across a population of genes ^4^. Gene molecules exhibit a range of promoter nucleosome configurations in activating conditions, including the fully nucleosomal promoter, the predominant configuration under repressing conditions ^5^. The structural variation in activating conditions arises locally and is explained by the assumption of a stationary stochastic process of nucleosome removal and reformation ^6,7^.

Both nucleosome formation and removal are catalyzed by ATP-dependent chromatin remodelers ^8,9^. In steady state, the net results appears to be no other than ATP hydrolysis, a futile cylce. While the purpose of nucleosome removal for transcription is evident, the purpose of promoter nucleosome reformation in activating conditions is not.

Transcription is controlled by transcriptional activators, which bind to specific sequence motifs in the promoter of selected genes and promote transcription by recruiting auxiliary factors to the promoter ^10^, including chromatin remodelers ^11^. Regulatory specificity or activator fidelity refers to the ability of the activator to stimulate transcription of its target gene compared to an off-target gene ^12^. While on-rates are closely similar between different DNA binding sites for the same eukaryotic activator, off-rates may range over two orders of magnitude ^13^. Activator specificity, thus, is a function of the difference in off-rates between specific and nonspecific DNA sequences.

Within the limits imposed by the energetics of activator-DNA binding, the specificity of sequence recognition may be increased by decreasing activator concentration ^12^. However, as activator concentration tends to zero, the search-time for specific binding sequences tends to infinity. Alternatively, the specificity of regulation may be improved by increasing the difference in standard free-energy change between correct and incorrect sequence binding reactions. This however poses the problem of how to turn genes quickly off once the activator is bound to its specific promoter sequence.

The problem of how to reconcile specificity with fast regulatory kinetics may be solved by kinetic proofreading of activator identity — the insertion of an activator-controlled, free energy-transmitting delay step into the activation path, which permits to kinetically distinguish between correct and incorrect promoter binding twice: before and after the delay step ^12,14^.

Activator-mediated ATP-dependent chromatin remodeling appears ideally suited for the kinetic proofreading of activator identity ^12^: Upon recruitment by promoter-bound activators, remodelers drive the promoter away from its thermodynamically preferred, repressed state; thus free energy is transmitted from the ATP-ADP work reservoir to the promoter ^14^. The stored energy is dissipated upon nucleosome reformation. The irreversible structural dynamics of the promoter drive both activator binding reactions, with the repressed and derepressed promoter, away from equilibrium: Activator-mediated recruitment of ATP-dependent chromatin remodelers and subsequent nucleosome removal continually removes the activator-bound promoter from the binding reaction between the repressed promoter and activator (*cf*. Fig. 1A, transition 2 → 3); consequentially, activator and repressed promoter associate faster than they dissociate. Likewise, the thermodynamically favored reformation of nucleosomes continually removes the activator-free promoter from the binding reaction between activator and derepressed promoter (*cf*. Fig. 1A, transition 4 → 1); activator and derepressed promoter dissociate faster than they unite. The activator, thus, preferentially binds the repressed rather than derepressed promoter and dissociates from the derepressed rather than repressed promoter (Fig. 1A). In this way, activator identity is appraised twice, before and after nucleosome removal.

**Fig 1.**
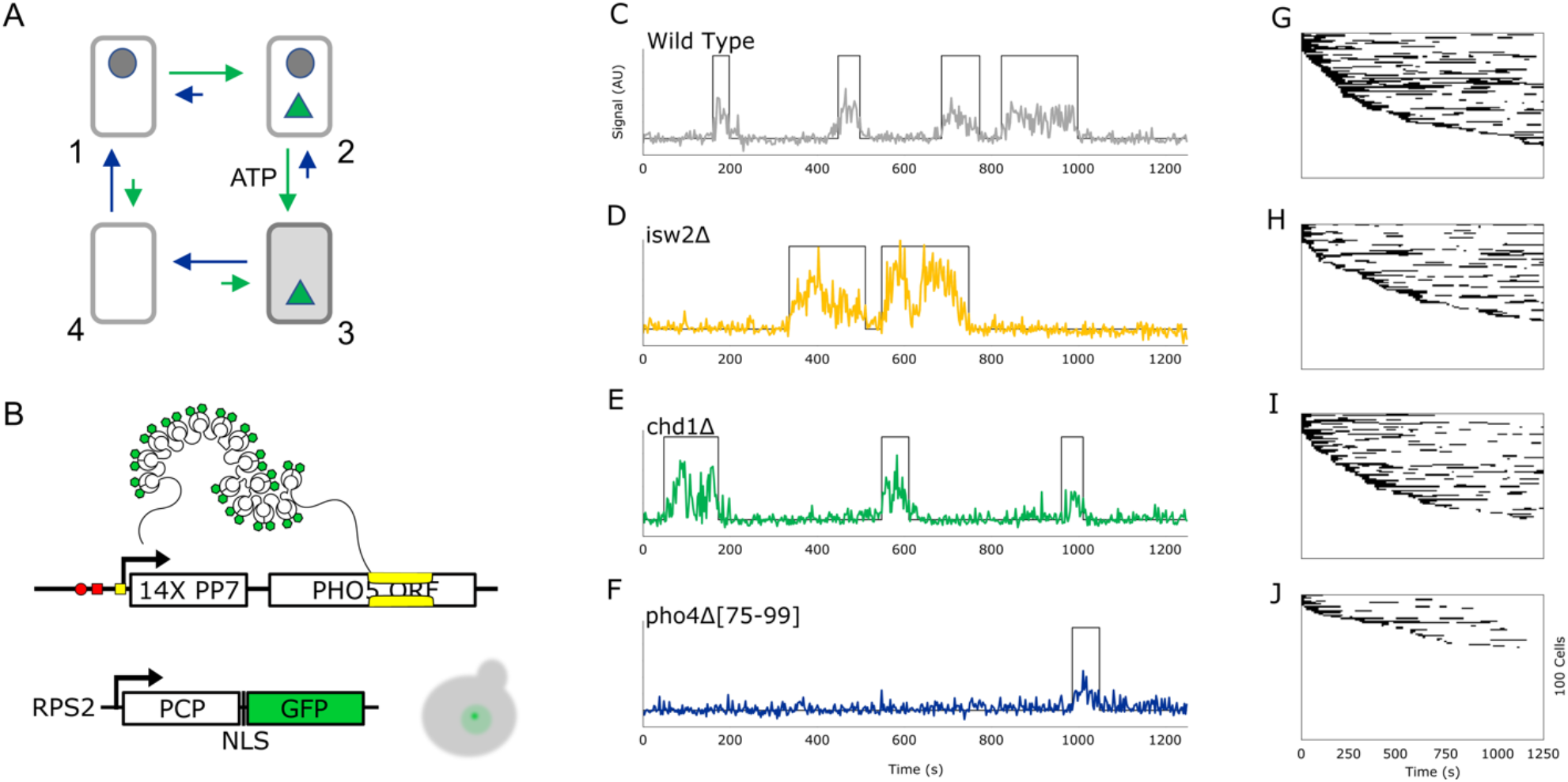
(A) Directed graph for the kinetic proofreading of activator identity ^12^. The promoter is depicted by a box, the bound transcriptional activator by a green triangle, a repressor whose removal (delay step) is required for transcription by a gray dot. Arrows (directed edges) indicate allowed transitions between microstates. Different arrow lengths indicate that clockwise transitions are always faster than counterclockwise transitions due to the transfer of free energy to the system in the transition from repressed to derepressed promoter in the presence of the activator, and free energy dissipation in the transition to the repressed state in the absence of the activator. The transcriptionally active state is highlighted in light gray. (B) The *PHO5* gene was modified by the splicing of a 14- mer cluster of binding sequences for the PP7 coat protein (PCP). PCP was expressed as a fusion with green fluorescent protein (GSP) with nuclear localization signal (NLS) under control of the *RPS2* promoter. The upstream activation sequences of *PHO5* are indicated by a red dot (UASp1) and red square (UASp2), the TATA box as a yellow square, the transcription start sites by bent arrows. The cartoon of a budding cells (in gray) shows that nuclear GFP is concentrated at the site of *PHO5* transcription (“transcription signal”). (C-F) Examples of single cell fluorescence sample paths for “wild type” (gray), *isw2*Δ (yellow), *chd1*Δ (green), and *pho4*Δ[75-90] (blue) cells. Based on phosphatase assays, the three mutations reduced *PHO5* expression to 50% (*isw2*Δ), 75% (*chd1*Δ), and 25% (*pho4*) of wild type expression. (G-J) Binary signal paths with black for transcription and white for no transcription of 100 cells for wild type (G), *isw2*Δ (H), *chd1*Δ (I), and *pho4*Δ[75-90] (J).

The resulting increase in activator fidelity, which is attained without increasing the affinity between activator and target promoter, allows for both fast regulatory kinetics (fast activator on- and off-rates) and fidelities that may breach the Hopfield fidelity barrier ^15^, the theoretical limit to regulatory specificity imposed by the energetics of activator-DNA binding for processes in equilibrium ^12,14^. In the sequel, we will refer to this conjecture simply as “proofreading hypothesis”.

We emphasize that the activator binding reactions are irreversible (i.e., out of equilibrium) not because the activator is actively removed from the DNA but because the promoter structure changes underneath the activator; the promoter states between activator binding and dissociation tend to be different (Fig. 1A). This temporal ordering of molecular events requires the dissipation of free energy ^14^.

Although the eukaryotic cell employs multiple free energy-consuming reactions to regulate transcription, it is not known whether the energy is expended to maintain the system away from equilibrium — as required for kinetic proofreading — or, alternatively, to overcome energetic barriers to accelerate the approach to equilibrium ^16^. Many aspects of transcriptional regulation are well explained on the assumption of a regulatory process in equilibrium. For instance, the random telegraph model of transcription, which has been invoked to explain higher than expected noise in gene expression, assumes that transcriptionally induced genes randomly toggle between two states, a transcriptionally conducive “ON” state and inconducive “OFF” state ^17^. In steady state, this process, like any process without closed loops (cycles), is also in equilibrium: forward and revers transitions between ON and OFF occur with equal speed, the process is said to be reversible or in detailed balance ^18^. With appropriate choice of rate constants, the random telegraph model was found to account very well for the statistical distribution of cytoplasmic transcripts across cells ^19^.

Recent technological advances have made possible the microscopic observation of transcriptional activity at single gene loci in living cells ^20^. However, while the perception of directionality evidently requires the observation of three or more states, microscopic analyses remain limited to the observation of transcription and its absence. With this limitation, equilibrium models are invoked by default. How to distinguish between reversible (in equilibrium) and irreversible (out of equilibrium) control processes is not evident.

## Results

We observed transcription at single *PHO5* gene copies of yeast — a classic paradigm for analysis of the relationship between chromatin and transcription ^21^ — by high-resolution multifocus fluorescence microscopy (MFM), which allows for the simultaneous acquisition of images at multiple focal planes ^22^.

Transcriptional activation of *PHO5* requires binding of the basic helix-loop-helix factor Pho4 to its upstream activation sequence UASp1 ^23^. Pho4 binding increases the probability of promoter states with fewer nucleosomes ^5^ — presumably by recruitment of ATP-dependent chromatin remodelers ^9,24^ — and thus allows Pho4 to access a second, previously occluded, binding site ^25,26^, UASp2, which accounts for about 75% of promoter activity ^27^. At high media concentrations of orthophosphate, Pho4 resides in the cytoplasm; its target genes, therefore, remain inactive. In cells that lack Pho80, a repressor of the PHO signaling pathway, Pho4 resides in the nucleus at all times. Hence, *pho80*Δ cells express *PHO5* constitutively, *i*.*e*., independently of environmental orthophosphate concentration.

We imaged *pho80*Δ cells, which induce *PHO5* constitutively ^28^. For simplicity, we will refer to the *pho80*Δ mutant as “wild type”. For microscopic observation, we labeled *PHO5* transcripts by inserting a cluster of 14 recognition-sequences for the RNA-binding coat protein of the PP7 phage into the 5′-untranslated region of the *PHO5* (Fig. 1B) ^20^. The coat protein was constitutively expressed as a fusion with green fluorescent protein (Fig. 1B). We acquired focal stacks, seven images spaced 500 *nm* apart along the optical axis, every 2.5 *s* over a period of 20 minutes (Fig. 1C-F).

We analyzed fluorescent time series by change point detection analysis ^29^ to segment time series into periods of transcriptional activity (ON) and periods of inactivity (OFF) (Figs. 1C-J), which allowed us to obtain statistical distributions for the lifetime of observed ON and OFF periods (Fig. 2).

**Fig. 2.**
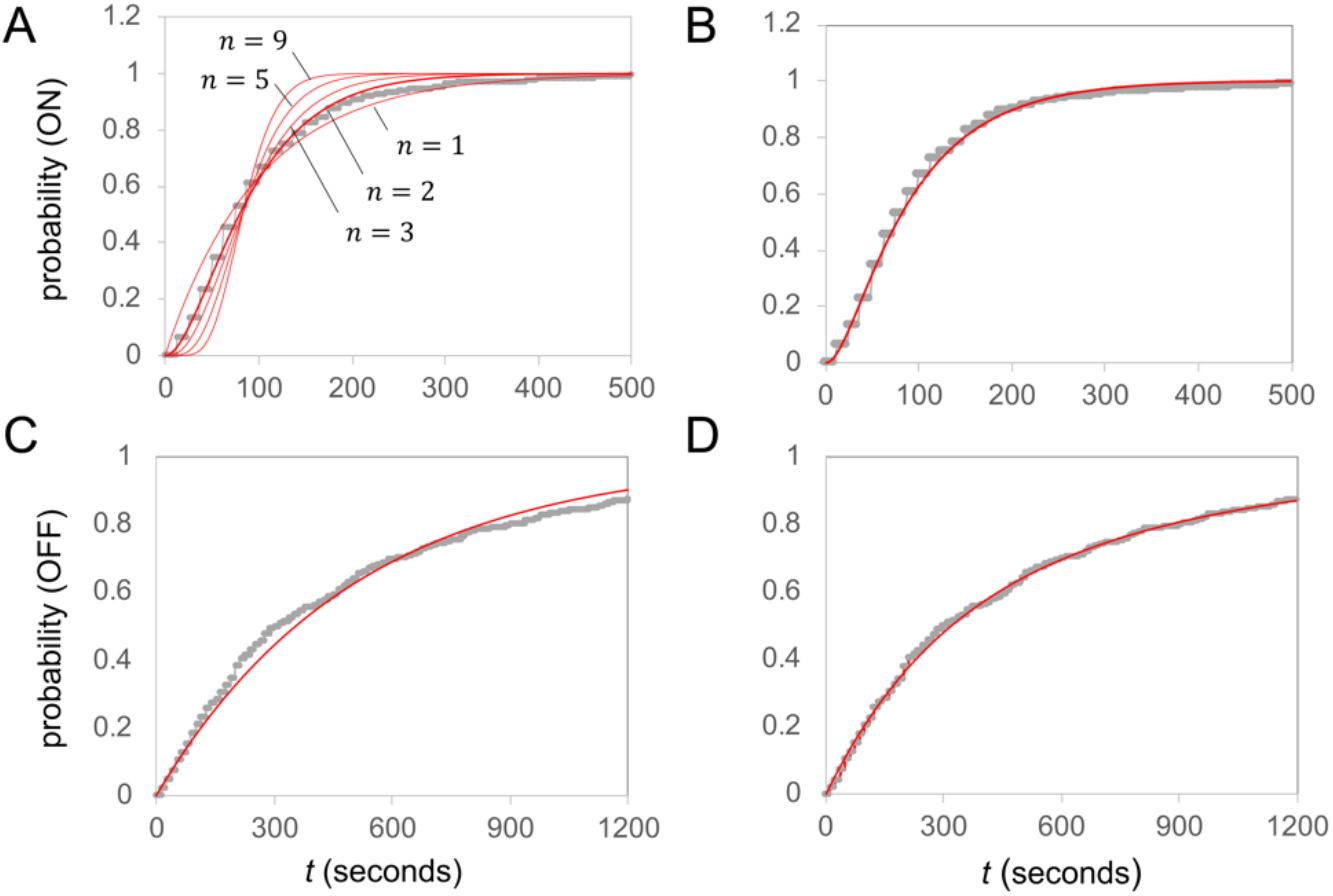
Distributions of process dwell times, *i*.*e*., the probability, *F*(*t*), that the dwell time in the observed state was ≤ *t*. (A) Experimental dwell time distribution in observed ON state is plotted in gray and best fit gamma distributions, 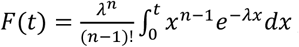, with shape parameter *n* = 1, 2, 3, 5 and 9 in red; *λ* > 0. (B) Same as A, but with best fit of 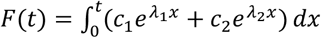, in red; *λ*_1_, *λ*_2_ < 0. (C) Experimental dwell time distribution in observed OFF state is plotted in gray, the best fit gamma distribution with *n* = 1 in red. (D) Same as C, but with two-exponential distribution (red) fit to the data (gray).

Setting the boundaries between ON and OFF periods, whether algorithmically or manually, is never free of arbitrary decisions. The algorithmic approach, however, has the advantage of articulating those decisions a priori, *i*.*e*., independently of the data. The random transitioning between ON and OFF periods engenders correlations between the transcription signals separated by fixed time intervals or “lag times”. The autocorrelation function of a stationary stochastic process gives this temporal correlation as a function of lag time. We calculated the autocorrelation function (*acf*) from fluorescence sample paths and from binary sample paths; the latter were obtained from the former by segmentation and setting the fluorescence signal of ON and OFF periods to 1 and 0, respectively (cf. Figs. 1G-J). The *acf* of binary sample paths is highly sensitive to the average length of ON and OFF periods (*cf*. SI). Yet, the *acfs* from fluorescence and corresponding binary sample paths were closely similar (*cf*. SI, Fig. S2), suggesting that the change point detection algorithm set boundaries more correctly, at least on average.

Due to the release of pyrophosphate and its subsequent cleavage into orthophosphate, nucleotide addition steps during transcription are effectively irreversible. Transcript elongation may be modeled, therefore, as a Poisson process. The time until the *n*^*th*^ nucleotide is added is then given by the sum of *n* independent, exponentially distributed random variables with a common mean. The sum is gamma distributed with integral shape parameter *n* ^30^. Thus, if ON periods corresponded to single transcription events, the length of ON periods would be gamma distributed with a large shape parameter. This, however, was not the case. With increasing shape parameter, the gamma distribution quickly tends toward a step function (Fig. 2A); and its density (the derivative of the distribution function) approaches a Dirac delta function. The best, although not good, fit to the data was obtained with a small shape parameter, *n* = 2 (*cf*. Fig. 2A).

The much wider than expected density functions may be explained on the assumption that observed ON periods delineated bursts of transcription, *i*.*e*., multiple transcript initiation events in quick succession, where each initiation event within the lifetime of the preceding nascent transcript extends the length of the observed ON period by the temporal gap between the two events. Burst are followed by periods of inactivity. Initiation events, thus, are temporally correlated.

Correlated initiation events are generally attributed to an underlying regulatory process that randomly transitions between two promoter microstates (assumption of a random telegraph process), one conducive to transcription (ON state), the other not (OFF state), where dwell times in microstates are exponentially distributed (Markov assumption). Thus, longer observed ON periods are explained by longer dwell times of the regulatory process in the microscopic ON state; the observed process is a reflection of the underlying regulatory process of transcript initiation, rather than transcript elongation.

The bursting hypothesis also explains the absence of a minimal duration of observed ON periods, corresponding to the expected lifetime of the nascent transcript (Fig. 2A); given, say, a polymerization speed of 33 nucleotides per second ^31^ and transcript length of 1.4 kb, it takes 40 seconds to synthesize a transcript, the minimal duration of an observed ON period. However, if we only detected transcription sites with more than one nascent transcript, shorter ON periods would correspond to bursts with fewer initiation events, the result of shorter dwell times in the ON state. For instance, if the gene returned to an OFF state after initiation of the *k*^*th*^ nascent transcript, where *k* is the minimal number for the detection of transcription, the duration of the observed ON period was determined not by the lifetime of the nascent transcript but the distance between the burst’s leading RNA polymerase and the gene’s poly(A)-signal, which may be arbitrarily short.

Alternatively, short windows of elevated signal may have mistakenly be identified as ON periods. While the latter hypothesis implies that the gene was transcriptionally active less often than inferred, the hypothesis that we detected only bursts with at least *k* nascent transcripts implies that we underestimated the number of ON periods.

Our live-cell microscopy indicated that the *PHO5* gene was transcriptionally active only 15% of the time (in wild type). In contrast, previous estimates based on the analysis of intrinsic protein noise suggested that the promoter was active 60% of the time ^5^. We furthermore analyzed cells by single-molecule fluorescence in situ hybridization (FISH) with DNA probes against PP7 RNA sequences. Consistent with the conclusion from protein noise analysis, FISH images showed that a substantial fraction of cells, 45% (in wild type), bore a single fluorescent punctum in the nucleus that was at least twice as bright as the average cytoplasmic punctum (Fig. 3A, 3B), suggesting the simultaneous presence of at least two transcripts at or near the *PHO5* locus. The quantitative discrepancy between FISH and live cell microscopy, which was also manifest in a previous study ^19^, is explained by the assumption that live-cell microscopy was limited to the observation of bursts of transcription — *i*.*e*., isolated, single transcription events were not observed. Thus, ON periods shorter than the expected lifetime of the nascent transcript are not likely an artefact of the change point detection algorithm. The percentage of cells with a nuclear punctum three or more times brighter than the average cytoplasmic punctum approximately matched the fraction of time the gene was seen to be transcriptionally active by live-cell imaging (*cf*. Fig. 3B), suggesting that observed ON periods corresponded to bursts with three or more nascent transcripts. The cytoplasmic and nuclear transcript numbers were nearly uncorrelated (Fig. 3C), as expected if the lifetime of the nuclear transcript — including a possible retention time after synthesis — was much shorter than the lifetime of the cytoplasmic transcript.

**Fig. 3.**
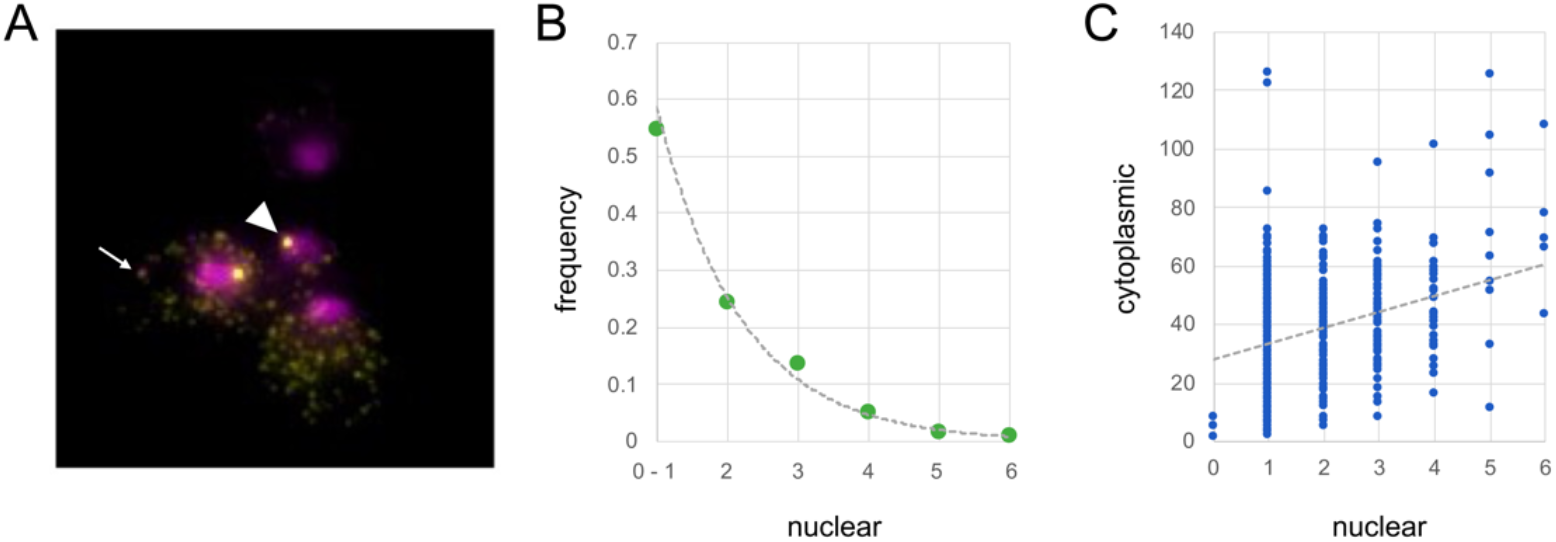
Number of cytoplasmic and nascent *PHO5* transcripts. *PHO5* transcripts were detected by FISH with DNA probes against PP7 sequences. (A) Fluorescence image. Nuclei were stained with DAPI (purple), *PHO5* transcripts are in yellow. A nuclear punctum that is thought to label the site of multiple nascent transcriptions is indicated by a white arrowhead. A white arrow points at a cytoplasmic dot, which is assumed to represent a single transcript. (B) Statistical frequencies of nuclear transcript numbers inferred from comparison of nuclear punctum intensities with mean cytoplasmic dot intensity; we analyzed 541 cells. Whether nuclear puncta with single transcript intensity corresponded to nascent transcripts is unknown; single transcript puncta could have lain above or below the nucleus and not within. (C) Scatter plot of nuclear versus cytoplasmic transcript numbers. With a Fano factor of 8.6 (variance over mean), the total noise of mRNA expression was markedly higher than expected for uncorrelated initiation events, Fano factor of 1.

Despite the complicated relationship between regulatory and observed process, multiple authors have reported exponentially distributed ON periods, suggesting a Markov process with one ON state (*e*.*g*., the random telegraph model). In contrast, we found that a close fit between observation and theoretical prediction required the assumption of two ON states, and not one; the exponential distribution (gamma distribution with shape parameter *n* = 1) did not fit the observed distribution of ON period-lengths (Fig. 2A).

In the case of two microscopic ON states, the distribution of the process dwell time in the compound ON state is given by 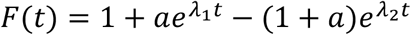 where *a* and *λ*_1_ < *λ*_2_ < 0 are constants (*cf*. SI). Hence, the corresponding density function, the derivative of *F*, is the sum of two weighted, decaying exponentials: 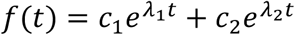. Remarkably, for all strains analyzed, we found that *c*_1_ + *c*_2_ = 0 (Fig. 4A-C; Table S1), which implies the existence of negative weights. With *f*(0) = *c*_1_ + *c*_2_ = 0, one obtains *c*_1_ = *λ*_1_*λ*_2_/(*λ*_1_ − *λ*_2_) < 0. As shown below, this finding has profound topological and thermodynamic implications.

**Fig. 4.**
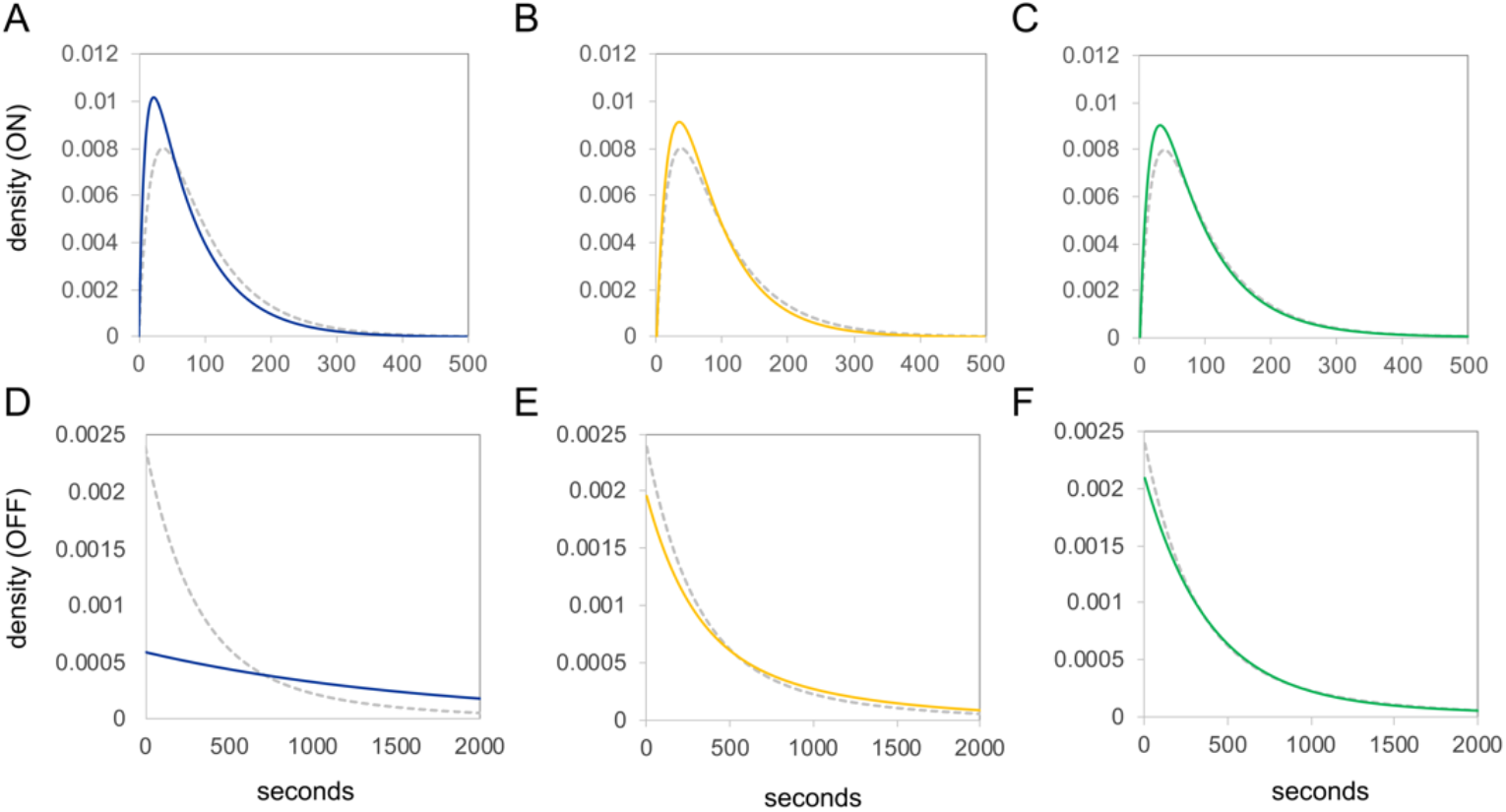
Probability density functions for the process dwell time in the observed ON state (A-C) and observed OFF state (D-F) in wild type (dashed gray), *pho4*Δ[75-90] cell (blue), *isw2*Δ cells (yellow), *chd1*Δ cells (green). Density functions were obtained by taking the derivative of the best fit two-exponential distribution function (*cf*. Fig. 2).

The process topology may be represented by a directed graph: a set of nodes, representing the accessible microstates of the process, and directed edges (arrows), indicating possible transitions between them.^32^ We assume that graphs are ergodic — *i*.*e*., every state can be reached from any other state by one or more transitions — or else the number of states may decrease as the process approaches steady state. For deriving the lifetime density function of the observed ON state, the observed OFF state may be represented by a single state (SI, Comment 3). We limit our analysis, therefore, to graphs with three states, two ON states and one OFF state (Fig. 5), although the latter may consist of several microstates.

**Fig. 5.**
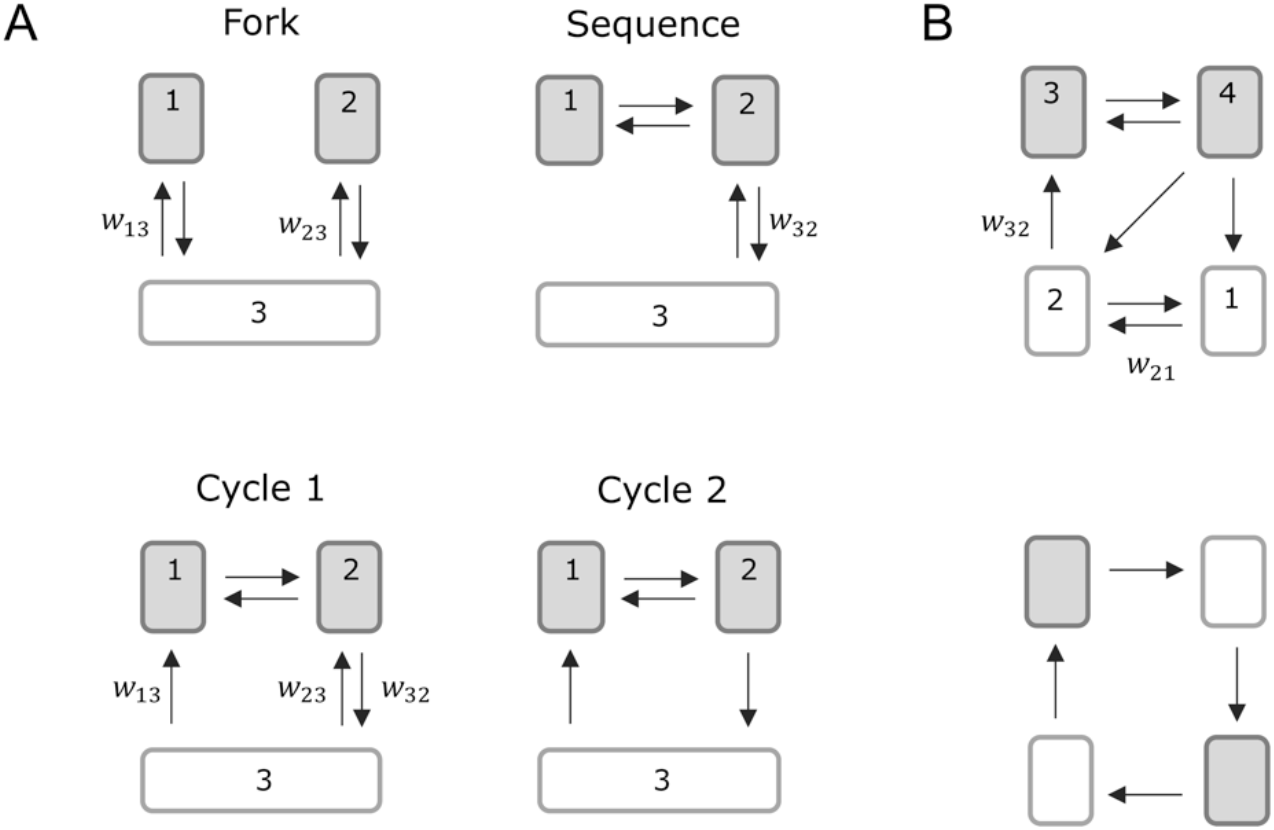
(A) Examples topological relationships of microscopic ON (OFF) states among each other (gray) and OFF (ON) states (white), which are presented as a single state. (B) Four-state graphs with two ON states (States 3 and 4) and two OFF states (States 1 and 2).

The topological relationship between the three states is described by either one of three kinds of graph (Fig. 5). The process may reach one microscopic ON state from the other only via passage through the OFF state (“Fork”). Alternatively, the process may visit both ON states in sequence but enters and exits the OFF state through only one of the two ON states and not the other (“Sequence”). The third kind encompasses all cyclic graphs, *i*.*e*., graphs with cycles. A cycle is a string of transitions whose initial and final state are the same without reversing itself at any point. Strongly connected graphs without cycles are called trees ^32^. Sequence and Fork are the only strongly connected graphs of their kind and together comprise all possible trees.

Inspection of the mathematical expressions for the density functions of the observed ON state shows that only cycles, and not trees, allow for negative exponential coefficient, *c*_*j*_ (SI, Theorems 2, 3, 4). Our experimental observations also rule out most cycles, since in general *c*_1_ + *c*_2_ ≠ 0 (cf. SI, Eq. [28]). Only cycles with irreversible entry into the observed ON state *via* one of its two microstates and irreversible exit *via* the other (Fig. 5, Cycle 2) fulfill *c*_1_ + *c*_2_ = 0 (*cf*. SI, Theorem 4).

By measurement of acidic phosphatase activity and single molecule FISH, we found that loss of ATP-dependent chromatin remodelers Chd1, Isw2, partial deletion of the Pho4 activation domain, and Swi2 decreased the steady state expression of *PHO5* to approximately 70%, 50%, 30% and 0% of wild type, respectively ^9,27^. The proofreading hypothesis posits that the activator controls burst frequency, in part, by recruitment of ATP-dependent chromatin remodelers (*cf*. Fig. 1A). Statistical analysis of observed ON and OFF period lengths corroborated this prediction; all three mutations reduced burst frequency; changes in gene expression were largely attributable to changes in burst frequency, with exception of the *isw2*Δ mutant (Table S2).

To test whether loss of ATP-dependent remodeling activity directly affected *PHO5* promoter chromatin, we isolated *PHO5* gene molecules from *chd1*Δ mutant cells and analyzed their nucleosome configuration by psoralen crosslinking and electron microscopy. Comparison with molecules isolated from wildtype cells ^5^ showed that the probability of nucleosome configurations with more nucleosomes increased, and the length of linkers spanning UASp1 decreased with loss of ATP-dependent chromatin remodeling activity (Fig. 6). Thus, the promoter more often assumed nucleosome configurations that prevail in repressing conditions ^5^, as expected on the assumption that the promoter continually returns to repressed states under activating conditions and that entry into bursts requires the expenditure of ATP; with fewer ATP molecules spent per time the promoter transitioned less often from OFF to ON.

**Fig. 6.**
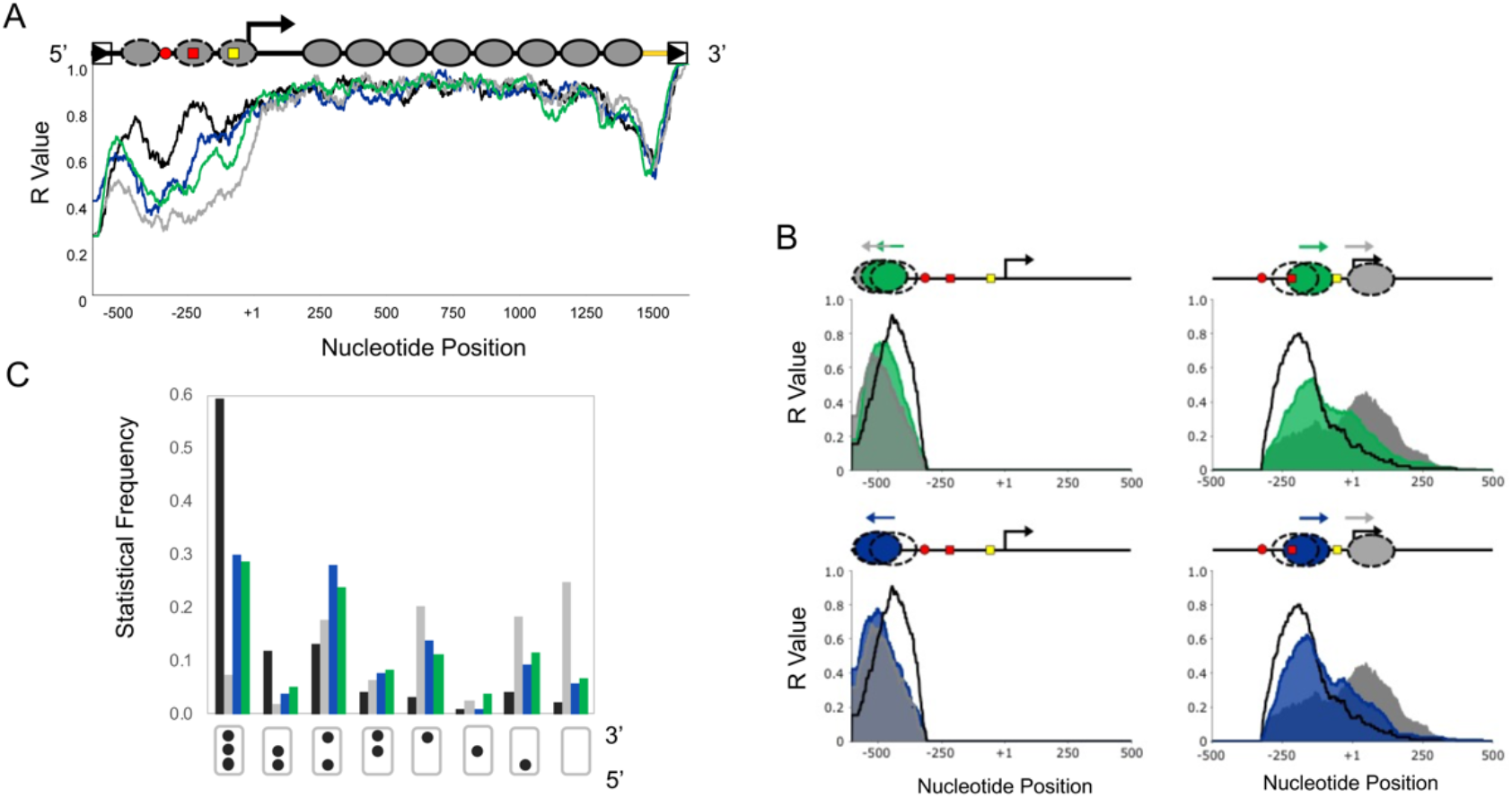
*PHO5* promoter nucleosome configurations from EM analysis of psoralen-crosslinked *PHO5* molecules. (A) Plot of R-value across the *PHO5* gene; the R value is the relative frequency of finding a nucleotide position single-stranded after psoralen-crosslinking of isolated *PHO5* chromatin rings and DNA denaturation. The R-value is an indicator of nucleosome occupancy, for psoralen crosslinks nucleosomal linker DNA and not core particle DNA. Gene molecules were analyzed by electron microscopy as previously described ^45^. A diagram of the *PHO5* gene is shown on top (*cf*. Fig. 1B). (B) For molecules with nucleosome-free UASp1(R value of zero) the R value of the adjacent 5’ and 3’ sequences is plotted against nucleotide position for *chd1*Δ pho80Δ (green) and *pho4*Δ[85-99] *pho80*Δ (blue). R values for the *pho80*Δ mutant (*PHO5* active) and *PHO80* (in high phosphate, *PHO5* inactive) are shown in gray and black, respectively. (C) Statistical frequencies of eight distinct nucleosome configurations in different strains: *PHO80* (black), *pho80*Δ (gray), *pho4*Δ[85-99] *pho80*Δ (blue) ^5^, and *chd1*Δ *pho80*Δ (green). The promoter is represented by a rectangle, occupied nucleosome positions by black dots. The 5’-terminal nucleosome is at the bottom, the 3’-terminal nucleosome at the top. Black: *PHO5* in repressing conditions (*PHO80* cells). Gray: *PHO5* in activating conditions (*pho80*Δ cells). Green: *PHO5* in *chd1*Δ *pho80*Δ cells. Blue: *PHO5* in *pho4*Δ[85-99] *pho80*Δ cells.

All three mutations also shortened burst length, albeit modestly (*cf*. Table S2). It is conceivable that this effect was caused by the decrease in burst frequency, for wider temporal gaps between bursts should reduce the chance of unresolved bursts. The length of observed ON periods, thus, is expected to become shorter at lower burst frequency. However, the comparison of autocorrelation functions from fluorescence sample paths also suggested a shortening of burst length in all mutant strains (*cf*. Fig. S1). Thus, changes in burst duration could not be attributed to unresolved bursts.

Two alternative, not mutually exclusive conjectures stand to reason that may explain shortened ON periods. First, the kinetic proofreading promoter-activator interactions requires that activators control both burst frequency and burst strength (the rate of transcript initiation during bursts) ^12^. A decrease in burst strength becomes manifest as shortened ON periods, for the chance of transcript initiation within the lifetime of the preceding nascent transcript decreases. Analysis of the activator mutation *pho4*Δ[75-90] bore out this expectation: The average length of ON periods decreased by approximately 20%, from 98 *s* to 81 *s* (*cf*. Table S2). Second, nucleosomes are thought to shorten the lifetime of activators on the DNA either by invading the activator binding site or by bending and twisting linker DNA upon interaction with each other ^33^. Thus, increased nucleosome occupancy and closer proximity of nucleosomes to activator binding sites is expected to decrease burst duration, provided the latter coincides with the lifetime of the activator on the DNA. Although loss of Chd1 activity led to a decrease in linker length and increase in promoter nucleosome occupancy similar to the *pho4*Δ[75-90] mutation (Figs. 6B, C), its effect on burst duration was markedly smaller (*cf*. Fig. 4A and C), suggesting that nucleosomes had little effect on the DNA-lifetime of the activator. Either specific activator-DNA interactions are able to resist nucleosomes, at least to some extent, or nucleosome transactions occur at a longer timescale compared to activator-promoter interactions; it is indeed difficult to see how else the specificity of transcriptional control could be maintained within a nucleosomal context.

The length distribution of observed OFF periods, like ON periods, fit the expectation of a Markov process with two microscopic OFF states, and not one (Fig. 2C, D) — the only exception was the *pho4* mutant, where the length of observed OFF periods was exponentially distributed, *i*.*e*., *c*_2_ = 0. Two OFF states suggest the existence of two rate-limiting steps along a path from transcriptional inactivity to bursting.

In contrast to observed ON periods, we found that *c*_1_, *c*_2_ ≥ 0 for all strains analyzed (Table S2). Thus, the topological relationships between microscopic OFF states with each other and ON states are potentially consistent with all graphs, except Cycle 2. The upper panel of Figure 5B depicts an example of a topological relationships between all four microstates — two ON states (States 1 and 2) and two OFF states (States 3 and 4) — that is permitted by our data; the topological relationships of ON states with each other and with OFF states are described by (are isomorphic to) Cycle 2; the relationships of OFF states with each other and with ON states are isomorphic to Cycle 1. The graph entails two paths out of the second ON state, consistent with the notion that both activator dissociation and nucleosome reformation may end a burst (*cf*. Fig. 1A). If the activator controls both the transition from one OFF state to the other (rate constant *w*_21_) and from the latter into one of the two ON states (rate constant *w*_32_), a mutation in the Pho4 activation domain would decrease both rate constants (Fig. 5B). In this case, *c*_2_ quickly approaches zero (SI, Theorem 5), which may explain the length distribution for observed OFF periods in the activator mutant. In contrast, the lower graph in Figure 5B is not permitted, although any process on this graph is irreversible, for the topological relationships of ON states (as well as OFF states) are isomorphic to the Fork.

## Discussion

Our analysis suggests that the regulatory process for transcription visits promoter microstates in a statistically preferred order, thus violating the detailed balance conditions for thermodynamic equilibrium. Refutation of the equilibrium hypothesis is the main result of our analysis.

This conclusion assumes that the observed process could be modeled by an irreversible process because the underlying regulatory process was irreversible. However, at the level of microscopic observation, the regulatory process was interwoven with the irreversible process of transcription. Could we have mistaken the latter for the first? We think the answer is no, if the conclusion is accepted that observed ON periods corresponded to bursts of transcription, trains of initiation events in quick succession whose observed length reflected the number of initiation events per train — and thus, albeit imperfectly, the dwell time of the process in microscopic states that are conducive to transcription — and not the lifetime of the nascent transcript. The latter possibility — that ON periods represented single transcription events — appears implausible, for it neither explains the wide distribution of ON period lengths nor the results from single molecule FISH. The quantitative discrepancy between both types of analysis, FISH and live cell microscopy, suggests furthermore that we only detected bursts of sufficient strength, and not isolated transcription events.

The possibility of missing transcription events in the observational record raises the question of whether transcriptional bursting was the result of incomplete observation: The gene might have been in an ON state at all times, but periods with sufficiently low density of transcription initiation were mistaken for OFF periods between bursts. At least two findings argue against this possibility. First, in the absence of bursting, mutations that decrease transcription must, on average, increase the temporal gaps between all transcription events. This would shorten the duration of apparent bursts. In contrast, we found that even mutations that severely reduced *PHO5* expression had comparatively little effect on the length distribution of observed ON periods (Fig. 4A); changes in expression were mostly explained by changes in burst frequency. Second, the Fano factor (variance over mean) of mRNA expression largely exceeded the expectation for a Poisson process (*i*.*e*., no bursting), nearly ten-fold (*cf*. legend to Fig. 3).

Given the complex relationship between observed and actual process, it is surprising that the length distribution of observed ON periods conformed closely to the expectation of a simple model: a Markov process with two ON states. Multiple ON states have been inferred before ^34,35^. What might those states be if indeed they physically exist? Eukaryotic promoters generally comprise multiple activator-binding sites. The *PHO5* promoter bears two Pho4 binding sites, UASp1 and UASp2. Transcription, therefore, may be driven by distinct patterns of activator binding site occupation, each representing a kinetically distinct microscopic ON state — e.g., our microscopic observations may have distinguished between ON states with one and two activator molecules bound at the promoter. Transitions out of microscopic ON states, either into OFF states or other ON states, thus correspond to single molecular events: the binding or dissociation of an activator to or from its target sequence. The lifetime of individual microscopic ON states is therefore exponentially distributed. This interpretation is in keeping with the proofreading hypothesis, which implies that bursts either end with the irreversible dissociation of activators from the derepressed promoter or reformation of a repressing nucleosome (Fig. 1A). We note that the average length of observed ON periods was closely similar to the average residence time of Pho4 at UASp2 in vitro, approximately 100 seconds ^13^, suggesting that transcription, at all times, requires the presence of an activator at the promoter.

Unidirectional transitions into and out of bursts (cf. Fig. 5, Cycle 2) violate the principle of microscopic reversibility — that any possible transition between two microscopic states is also possible in the opposite direction. However, with the dissipation of a sufficiently large amount of free energy along any microscopic path, the probability of the reverse path may become virtually impossible. The free energy change of ATP hydrolysis by cytoplasmic enzymes may be as large as −57 *kJ*/*mol* or −23 *k*_*B*_*T* per molecule ^36^. With the detailed fluctuation theorem of stochastic thermodynamics ^37^ follows, therefore, that the hydrolysis of one ATP molecule along any cyclic microscopic path reduces the probability of the reverse cycle relative to the forward cycle by a factor of *e*^23^ or 10^10^.

Our major analytical result — that *c*_1_ + *c*_2_ = 0 implies the existence of an irreversible process (*cf*. SI, Theorem 5) — is closely related to Tu’s theorem, which states that detailed balance prohibits dwell time densities with negative coefficients ^38^. Thus, *c*_1_ + *c*_2_ = 0 implies that the process is irreversible and cyclic if in steady state, for steady-state processes on trees are reversible (*i*.*e*., in equilibrium) ^18^. In contrast, we found that trees do not allow for negative *c*_*j*_′*s*, regardless of whether the process is in steady state (*cf*. SI, Theorem 2), and that *c*_1_ + *c*_2_ = 0 imposes a topology that is incompatible with detailed balance (*cf*. SI, Theorem 5). While our analysis was limited to observed states that comprise two microstates, Tu proved his theorem for any time-homogeneous Markov process ^38^.

We note that irreversible processes may appear reversible to an observer who insufficiently discriminates between microstates. As an example, we consider a steady-state stochastic process on the cyclic graph depicted in the upper panel of Fig. 5B. The inability to distinguish between microstates 1 and 2 on one hand, and microstates 3 and 4 on the other, converts the cyclic graph into a tree; the observed process becomes a random telegraph process, which in steady state is reversible, although the underlying microscopic process is not. This may explain why irreversible microscopic processes are difficult to detect and why equilibrium models may successfully be applied to irreversible processes.

The proofreading hypothesis proffers multiple predictions for testing. For proofreading, the promoter must continually fall back into a state that precedes one or more delay steps (Fig. 1A). This may explain why the promoter revisits repressing nucleosome configuration in activating conditions; promoter chromatin of transcriptionally active promoters is structurally heterogeneous ^5,27,39–41^. The regulatory process is devoid of memory; the system, thus, continually recapitulates, at least in part, the induction process. Transcription, therefore, occurs in stochastic bursts ^12,42^; activators control transcription primarily at the level of burst frequency (*cf*. Table S2) ^43^; and transcription in activating conditions requires the continual employment of chromatin remodelers (Fig. 6). Transcription initiation must be closely tied to the presence of activators on the DNA. Thus, burst length is limited by the DNA residence time of activators. All these expectations were born out by this and other analyses ^19,20,42^. Most importantly, the control process must be irreversible. In our analysis, we detected a telltale sign of irreversibility: peaked dwell time densities for observational states. Peaked dwell time densities have been found before in mammalian gene expression ^35,44^. In the absence of a theory that imparted significance to this finding, its thermodynamic and topological implications remained unnoted, however.

## Methods

### Strains

All strain for live cell fluorescent microscopy were derived from yM2.1 ^4^, which contains sequence insertions upstream and downstream of the *PHO5* gene for enzymatic release of the *PHO5* gene from its chromosomal locus. Insertion of the PP7 binding sequence clusters was achieved by homologous recombination; the cluster along with a selection marker (KanMX) flanked by loxP recombination sequences was PCR amplified from plasmid pTL31 using primers p446 (5’- CTTCATCTCTCATGAGAATAAGAACAACAACAAATAGAGCAAGCAAATTCGAGATTACCAcaaagtgggagcgaggagatcc-3’) and p447 (5’- AATGGTACCTGCATTGGCCAAAGAAGCGGCTAAAATTGAATAAACAACAGATTTAAACATgcataggccactagtggatctg-3’) whose 5’-ends are homologous to the *PHO5* 5’ UTR. The selective marker was removed by transformation with plasmid pSH47 and induction of the site-specific recombinase. The PP7 coat protein-GFP fusion was expressed under control of the *RPS2* promoter; the fusion gene was inserted at the *ura3* locus by homologous recombination with a DNA fragment generated by digestion of pTL205 (a derivative of pSIVURA3 ^46^) with PacI. The fragment bore the *URA3* gene of *Candida albicans* for selection. Activation of the *PHO* pathway was achieved by deletion of *PHO80* using transforming DNA pCM67 digested with EcoRI. Chromatin remodeler deletions were also generated by homologous recombination: *isw2* (pCM122 digested with XbaI and KpnI) and *chd1* (pCM123 digested with XbaI and KpnI). The activator mutants were generated by deletion of Pho4 and subsequent insertion of an altered copy. This was achieved by selection and counter selection of *URA3*. Deletion was accomplished with transforming DNA pCM4 digested with BamHI and SalI while the mutant (Pho4 amino acids 75-90 deleted) was returned with pCM64 also digested with BamHI and SalI. All strains for single molecule FISH were derived from YS18 (the parental strain of yM2.1). Single molecule FISH experiments were performed with derivatives of YS18 (which lacks sequences for circle formation).

### Live-Cell Imaging

Yeast Cells were grown in 50 ml of liquid culture to 3-5 × 10^7^ cells/ml. Cells were then concentrated and spotted onto an agar patch on a #1.5 coverslip, as described elsewhere ^47^. For coverslip preparation we used the mold described by Skinner *et al*., (2013). Prior to imaging, cells were then incubated at 30°*C* for 30 minutes. A stage top incubator (OkoLab) was used to maintain a constant temperature of 30°*C* during imaging. MFM images were recorded with on an EMCCD camera (Andor Ixon Ultra). Images were taken every 2.5 second over 20 minutes with an exposure time of 250 *ms* and a laser power of 30% (Lambda XL). Excitation and emission paths were filtered using a Semrock GFP-30LP-B Brightline filter set. The MFM was assembled as previously described ^22,49^ and appended to the light path of an existing inverted widefield microscopy chassis (Nikon Eclipse TI), which maintained a 60 × objective (plano Apo 60X/1.4 oil immersion).

Fluctuation analyses were performed in four steps: (*i*) Identification of transcription sites. (*ii*) Assignment of transcription sites to nuclei. (*iii*) Tracking of each transcription sites over time. (*iv*) Quantification of signal intensity at transcription site.

To identify transcription sites, we first calculated a maximum projection of the z-stack for each time point; the maximum projection minimizes fluctuations in puncta intensity due to RNA movement in the z-direction (*i*.*e*., along the optical axis) during imaging. We then applied a band pass filter to maximum projections with a bandwidth to match the width of the point spread function of the microscope (approximately 1.5 pixels); the band pass filter reduces the rate of false positives due to background nuclear fluorescence caused by cytoplasmic RNA and unbound coat protein ^50^. To find local maxima (*i*.*e*., candidate transcription sites) we applied a search window of 4 pixels and a threshold of five standard deviations above the mean filtered image signal. We assigned transcription sites to nuclei using the background nuclear fluorescence to determine nuclear boundaries. We always identified the brightest site within a nuclear boundary as “the site of transcription”. Time series were generated by following individual transcription sites over time. When no candidate transcription site could be detected within a nucleus at a given time point, the location of the previously identified transcription site was used instead. In the case of no previous transcription site, we used the location of the brightest nuclear pixel in instead. Finally, to quantify the signal intensity of transcription sites, we used a Gaussian mask algorithm ^50^.

We corrected for photobleaching by detrending time series; a trend was inferred by regression analysis of the “average” time series, as described by Wu *et al*. (2007). Variation in nuclear intensity creates variation in background intensity. To correct for offsets of time series against each other along the y-axis (signal intensity), we subtracted from each time point the mode of the estimated probability function (kernel density estimate) of signal intensity of each time series. The mode was used because, unlike mean and median, it correctly reflects the background on the assumption that the most likely number of nascent transcripts is zero — an assumption borne out by our observations.

We assumed that sample paths (i.e., fluorescence signal traces) were step functions subject to Gaussian noise. Steps, time intervals of signal that appeared to be drawn from the same Gaussian distribution, were determined by change point detection or CPD analysis ^31^; we used a window-based search method together with a Gaussian cost function in the “ruptures” python package ^31^. We observed essentially two types of steps: low-noise steps at lower signal (OFF steps) and high-noise steps at higher signal (ON steps). CPD hyperparameters (e.g., window size) were selected to reduce the number of sequential low-noise steps, on the assumption that the background signal — the signal during OFF periods — is step-free. In contradistinction, the transcription signal — the signal during ON periods — may encompass multiple steps, corresponding to different numbers of nascent transcript.

### Model Fitting

Mathematical functions were fit to experimental distributions using the FindFit function in Mathematica 9. For distribution functions with two exponentials,

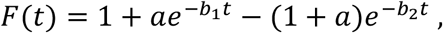

the search for the parameters *a, b*_1_, *b*_2_ was constrained by putting d*F*(0)/*dt* = −*ab*_1_ + (1 + *a*)*b*_2_ ≥ 0 to avoid negative probabilities.

### Autocorrelation Analysis

For individual sample paths, the autocorrelation at lag-time *k* was calculated according to

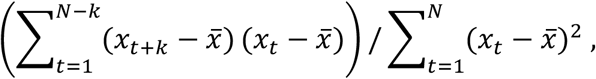

where *x*_*t*_ is the signal intensity of the transcription site (see above) at time point *t*, 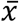 is the global mean — obtained by averaging over all time points of all time-series, as suggested by Coulon and Larson ^52^ — and *N* is the total number of time points per time series ^53^. Experimental autocorrelation functions were then obtained by averaging autocorrelations at lag-time *k* over all recorded sample paths.

### Single Molecule FISH

Yeast cells were grown in liquid culture (50 *ml*) to mid log-phase, crosslinked with formaldehyde, lysed with lyticase and adhered to poly-L-lysine coated coverslips, as previously described ^54,55^. Coverslips were hybridized, for 5 hours at 37°C, with 2.5 *nM* of fluorescently labeled antisense-DNA probe against PP7 binding sequences. Probes were labeled with Quasar 570 and Quasar 670 (Biosearch Technologies). For microscopy, coverslips were mounted onto glass slides with mounting media containing DAPI (4’, 6-diamidino-2-phenylindole; ProLong Gold, Life Technologies).

Cytoplasmic and nascent RNA were identified by using background fluorescence and DAPI staining to estimate cellular and nuclear boundaries, respectively. The number of nascent RNAs was calculated by normalizing the transcription site intensity with the average intensity of cytoplasmic transcripts. All analysis was accomplished using custom-made scripts and previously published code ^52^.

### PHO5 Isolation and EM Analysis

Gene isolation and psoralen-EM analysis were performed as previously described ^45^. Molecules were analyzed with custom-made programs written in Python. At least 200 *PHO5* molecules were analyzed for each strain.

## Supporting information

Mathematical background and Supplemental Tables

## Acknowledgements

This work was support by the National Science Foundation, Award #: 2111763 to H.B.

## Citations

1. Thastrom, A. et al. Sequence motifs and free energies of selected natural and non-natural nucleosome positioning DNA sequences. J Mol Biol 288, 213–229 (1999).

2. Kornberg, R. D. & Lorch, Y. Primary Role of the Nucleosome. Molecular Cell (2020) doi:10.1016/j.molcel.2020.07.020.

3. Almer, A., Rudolph, H., Hinnen, A. & Hörz, W. Removal of positioned nucleosomes from the yeast PHO5 promoter upon PHO5 induction releases additional upstream activating DNA elements. EMBO J. 5, 2689–2696 (1986).

4. Boeger, H., Griesenbeck, J., Strattan, J. S. & Kornberg, R. D. Nucleosomes unfold completely at a transcriptionally active promoter. Mol. Cell 11, 1587–1598 (2003).

5. Brown, C. R., Mao, C., Falkovskaia, E., Jurica, M. S. & Boeger, H. Linking stochastic fluctuations in chromatin structure and gene expression. PLoS Biol 11, e1001621 (2013).

6. Boeger, H. Nucleosomes, transcription, and probability. Mol Biol Cell 25, 3451–3455 (2014).

7. Brown, C. R. & Boeger, H. Nucleosomal promoter variation generates gene expression noise. Proc Natl Acad Sci U S A 111, 17893–17898 (2014).

8. Clapier, C. R. & Cairns, B. R. The biology of chromatin remodeling complexes. Annu Rev Biochem 78, 273–304 (2009).

9. Brown, C. R., Mao, C., Falkovskaia, E., Law, J. K. & Boeger, H. In Vivo role for the chromatin-remodeling enzyme SWI/SNF in the removal of promoter nucleosomes by disassembly rather than sliding. J. Biol. Chem. 286, 40556–40565 (2011).

10. Ptashne, M. How eukaryotic transcriptional activators work. Nature 335, 683–689 (1988).

11. Cosma, M. P., Tanaka, T. & Nasmyth, K. Ordered recruitment of transcription and chromatin remodeling factors to a cell cycle-and developmentally regulated promoter. Cell 97, 299–311 (1999).

12. Shelansky, R. & Boeger, H. Nucleosomal proofreading of activator–promoter interactions. Proc. Natl. Acad. Sci. U. S. A. 117, 2456–2461 (2020).

13. Maerkl, S. J. & Quake, S. R. A systems approach to measuring the binding energy landscapes of transcription factors. Science (80-.). 315, 233–237 (2007).

14. Boeger, H. Kinetic Proofreading. Annu. Rev. Biochem. (2022) doi:10.1146/annurev-biochem-040320-103630.

15. Estrada, J., Wong, F., DePace, A. & Gunawardena, J. Information Integration and Energy Expenditure in Gene Regulation. Cell 166, 234–244 (2016).

16. Wong, F. & Gunawardena, J. Gene Regulation in and out of Equilibrium. Annu. Rev. Biophys. 49, 199–226 (2020).

17. Ko, M. S. A stochastic model for gene induction. J Theor Biol 153, 181–194 (1991).

18. Kelly, F. P. Reversibility and stochastic networks. (John Wiley and Sons Ltd, 1979).

19. Donovan, B. T. et al. Live-cell imaging reveals the interplay between transcription factors, nucleosomes, and bursting. EMBO J. (2019) doi:10.15252/embj.2018100809.

20. Larson, D. R., Zenklusen, D., Wu, B., Chao, J. A. & Singer, R. H. Real-time observation of transcription initiation and elongation on an endogenous yeast gene. Science (80-.). 332, 475–478 (2011).

21. Korber, P. & Barbaric, S. The yeast PHO5 promoter: from single locus to systems biology of a paradigm for gene regulation through chromatin. Nucleic Acids Res 42, 10888–10902 (2014).

22. Abrahamsson, S. et al. Fast multicolor 3D imaging using aberration-corrected multifocus microscopy. Nat Methods 10, 60–63 (2013).

23. Vogel, K., Horz, W. & Hinnen, A. The two positively acting regulatory proteins PHO2 and PHO4 physically interact with PHO5 upstream activation regions. Mol Cell Biol 9, 2050–2057 (1989).

24. Barbaric, S. et al. Redundancy of chromatin remodeling pathways for the induction of the yeast PHO5 promoter in vivo. J Biol Chem 282, 27610–27621 (2007).

25. Mao, C., Brown, C. R., Griesenbeck, J. & Boeger, H. Occlusion of regulatory sequences by promoter nucleosomes In Vivo. PLoS One 6, 2–11 (2011).

26. Venter, U., Svaren, J., Schmitz, J., Schmid, A. & Horz, W. A nucleosome precludes binding of the transcription factor Pho4 in vivo to a critical target site in the PHO5 promoter. Embo J 13, 4848–55. (1994).

27. Mao, C. et al. Quantitative analysis of the transcription control mechanism. Mol. Syst. Biol. 6, 1–12 (2010).

28. O’Neill, E. M., Kaffman, A., Jolly, E. R. & O’Shea, E. K. Regulation of PHO4 nuclear localization by the PHO80-PHO85 cyclin-CDK complex. Science (80-.). 271, 209–212 (1996).

29. Truong, C., Oudre, L. & Vayatis, N. Selective review of offline change point detection methods. Signal Processing 167, 107299 (2020).

30. Feller. An introduction to probability theory and its applications. Volume II. (John Wildey & Sons, Inc., 1971).

31. Mason, P. B. & Struhl, K. Distinction and relationship between elongation rate and processivity of RNA polymerase II in vivo. Mol. Cell 17, 831–840 (2005).

32. Gunawardena, J. A linear framework for time-scale separation in nonlinear biochemical systems. PLoS One 7, (2012).

33. Fierz, B. Dynamic Chromatin Regulation from a Single Molecule Perspective. ACS Chem Biol 11, 609–620 (2016).

34. Corrigan, A. M., Tunnacliffe, E., Cannon, D. & Chubb, J. R. A continuum model of transcriptional bursting. Elife 5, 1–38 (2016).

35. Tantale, K. et al. A single-molecule view of transcription reveals convoys of RNA polymerases and multi-scale bursting. Nat. Commun. 7, 12248 (2016).

36. Nicholls, D. G. & Ferguson, S. J. Bioenergetics 4. Academic ORess (Academic Press, 2013).

37. Broeck, C. Van Den & Esposito, M. Ensemble and trajectory thermodynamics : A brief introduction. Physica A 418, 6–16 (2015).

38. Tu, Y. The nonequilibrium mechanism for ultrasensitivity in a biological switch: Sensing by Maxwell’s demons. Proc. Natl. Acad. Sci. U. S. A. 105, 11737–11741 (2008).

39. De Bernardin, W., Koller, T. & Sogo, J. M. Structure of in-vivo transcribing chromatin as studied in simian virus 40 minichromosomes. J Mol Biol 191, 469–482 (1986).

40. Boeger, H., Griesenbeck, J. & Kornberg, R. D. Nucleosome Retention and the Stochastic Nature of Promoter Chromatin Remodeling for Transcription. Cell 133, (2008).

41. Stergachis, A. B., Debo, B. M., Haugen, E., Churchman, L. S. & Stamatoyannopoulos, J. A. Single-molecule regulatory architectures captured by chromatin fiber sequencing. Science (80-.). (2020) doi:10.1126/science.aaz1646.

42. Rodriguez, J. & Larson, D. R. Transcription in Living Cells: Molecular Mechanisms of Bursting. Annu. Rev. Biochem. 89, 189–212 (2020).

43. Fukaya, T., Lim, B. & Levine, M. Enhancer Control of Transcriptional Bursting. Cell 166, 358–368 (2016).

44. Suter, D. M. et al. Mammalian genes are transcribed with widely different bursting kinetics. Science (80-.). 332, 472–474 (2011).

45. Brown, C. R. et al. Chromatin structure analysis of single gene molecules by psoralen cross-linking and electron microscopy. Methods Mol. Biol. 1228, 93–121 (2015).

46. Wosika, V. et al. New families of single integration vectors and gene tagging plasmids for genetic manipulations in budding yeast. Mol. Genet. Genomics 291, 2231–2240 (2016).

47. Zawadzki, K. & Broach, J. A rapid technique for the visualization of live immobilized yeast cells. J. Vis. Exp. (2006) doi:10.3791/84.

48. Skinner, S. O., Sepúlveda, L. A., Xu, H. & Golding, I. Measuring mRNA copy number in individual Escherichia coli cells using single-molecule fluorescent in situ hybridization. Nat. Protoc. (2013) doi:10.1038/nprot.2013.066.

49. Abrahamsson, S. et al. Multifocus microscopy with precise color multi-phase diffractive optics applied in functional neuronal imaging. Biomed Opt Express 7, 855–869 (2016).

50. Thompson, R. E., Larson, D. R. & Webb, W. W. Precise nanometer localization analysis for individual fluorescent probes. Biophys. J. (2002) doi:10.1016/S0006-3495(02)75618-X.

51. Wu, Z., Huang, N. E., Long, S. R. & Peng, C. K. On the trend, detrending, and variability of nonlinear and nonstationary time series. Proc. Natl. Acad. Sci. U. S. A. (2007) doi:10.1073/pnas.0701020104.

52. Coulon, A. & Larson, D. R. Fluctuation Analysis Dissecting Transcriptional Kinetics with Signal Theory. Methods Enzymol. 572, 159–191 (2016).

53. Percival, D. B. & Walden, A. T. Spectral analysis for physical applications. (Cambridge University Press, 1993).

54. Trcek, T. et al. Single-mRNA counting using fluorescent in situ hybridization in budding yeast. Nat. Protoc. (2012) doi:10.1038/nprot.2011.451.

55. Patel, H. P., Brouwer, I. & Lenstra, T. L. Optimized protocol for single-molecule RNA FISH to visualize gene expression in S. cerevisiae. STAR Protoc. 2, 100647 (2021).

56. Cinlar, E. Introduction to stochastic processes. (Dover Publications, 2013).

57. Mirzaev, I. & Gunawardena, J. Laplacian Dynamics on General Graphs. Bull. Math. Biol. 75, 2118–2149 (2013).

58. D. Colquhoun and A. G. Hawkes Source. On the Stochastic Properties of Single Ion Channels. Proc. R. Soc. London. Ser. B, Biol. Sci. 211, 205–235 (1981).

59. Grimmet, G. & Stirzaker, D. Probability and random processes. (Oxford university press, 2001).

