## Supplementary material for "A Telltale Sign of Irreversibility in Transcriptional Regulation": Mathematical background and Supplemental Tables

#### SUPPLEMENTAL INFORMATION

***Theoretical background and justification are given for mathematical results used in the main text — most importantly, Theorems 1 and 4 —followed by a brief discussion of the autocorrelation function. Tables S1 and S2 contain parameters obtained from model fitting. Table S3 lists the dwell time density functions of the A-set for the directed graphs of Fig. 5B.***

#### Dwell Time Distributions

Let $Y = \{Y_t: \Omega \longrightarrow S | t \in \mathbb{R}_{\leq 0}\}$ be a time-homogeneous Markov process with finite state space $S$, which is partitioned into two subsets $A$ and $B$ (ON and OFF) with $S = A \cup B$. The holding or dwell time of the process in $A$ is a continuous random variable, $H_A$. What is the probability density function of $H_A$? The answer is given by Theorem 1 (see p. 6). In the following we lay out how to get there.

Let $n$ be the total number of states and $s < n$ the number of $A$ states. The number of $B$ states therefore is $n - s$. For time-homogeneous Markov processes, the rate of transition from state $i$ to $j$ is given by $w_{ji}p_i(t)$, where $p_i(t)$ is the probability that the process resides in state $i$ at time $t$ and $w_{ji}$ is a rate constant [1]. The column vector $\boldsymbol{p}(t) = (p_1(t), \ldots, p_n(t))^T$ is called the probability mass function at time $t$ [(1)]. Provided the process graph is strongly connected — *i.e.*, every state may be reached from any other state by one or more transitions — the process approaches a uniquely defined distribution (independent of the initial distribution). This distribution is called the stationary distribution $\boldsymbol{p}$ [2]. The process, then, is in steady state, *i.e.*, $dp_k(t)/dt = 0$ for all $k \in S$.

The "generator" of the process [1] is the square matrix

$$\mathrm{W} = \begin{pmatrix} w_{11} & \cdots & w_{1n} \\ \vdots & \ddots & \vdots \\ w_{n1} & \cdots & w_{nn} \end{pmatrix}$$

[(1)] The superscript $T$ indicates the transpose of the row vector (which therefore becomes a column vector).

where

$$w_{jj} \equiv -\sum_{k \neq j} w_{jk} \,. \qquad [1]$$

In words, the diagonal element $w_{jj}$ is the negative sum of the remaining elements in column $j$.

We enumerate the elements of $S$ such that states $1, \dots, s$ are $A$ states and states $s+1, \dots, n$ are $B$ states. Correspondingly, we partition the generator into the following sub-matrices [3]:

$$\mathrm{W}_{AA} = (w_{ji}) \text{ where } i, j \in A; \quad \mathrm{W}_{AB} = (w_{ji}) \text{ where } j \in A, i \in B; \ \textit{etc.}$$

The generator, then, is

$$\mathrm{W} = \begin{pmatrix} \mathrm{W}_{AA} & \mathrm{W}_{AB} \\ \mathrm{W}_{BA} & \mathrm{W}_{BB} \end{pmatrix}.$$

(Note that the diagonal elements of W are contained in the diagonal matrices $\mathrm{W}_{AA}$ and $\mathrm{W}_{BB}$.) Other vectors and matrices are partitioned accordingly — *e.g.*, $\boldsymbol{p} = (\boldsymbol{p}_A, \boldsymbol{p}_B)^T$.

The vectors $\boldsymbol{u}_A$ and $\boldsymbol{u}_B$ designate the row vectors $(1, \dots, 1)$ of length $s$ and $n - s$, respectively. For the probability that the process leaves $A$ no earlier than $t$ but no later than $t + h$, given that it started in $i \in A$ at $t = 0$ we write

$$P(t \leq H_A < t + h | Y_0 = i \in A) \,.$$

To find this probability, which is the integral over the density function of $H_A$ between $t$ and $t + h$, we follow Colquhoun and Hawkes and consider the *modified* process in which all $B$ states are absorbing [3]. Thus, $j \in B$ is reached from $i \in A$ by crossing the boundary between $A$ and $B$ no more than once (*i.e.*, $\mathrm{W}_{AB} = \mathbf{0}$), and probability mass does not disperse between $B$ states ($\mathrm{W}_{BB} = \mathbf{0}$).([2])

For the *modified* process, let

$$F_{ji}(t) = P(Y_t = j | Y_0 = i)$$

([2]) Here and in the following, $\mathbf{0}$ is the zero matrix; its dimensions will be clear from the context, including the zero row and column vector.

be the probability that the process is in state $j$ at time $t$, given that the process was in state $i$ at time 0 for all $i, j \in S$. Furthermore, let $F = (F_{ji})$ be the matrix of all "transition functions", $F_{ji}$. In the following, let $j \in B$ and $i \in A$. With Chapman-Kolmogorov [1], we may write

$$F_{ji}(t+h) = \sum_{r \in S} F_{jr}(h) F_{ri}(t)\,. \qquad [2]$$

However, since $B$ states are absorbing, $F_{jr}(h) = 0$ for all $r \in B$ with $r \neq j$, and $F_{jr}(h) = 1$ for $r = j$. Hence,

$$F_{ji}(t+h) = \sum_{r \in A} F_{jr}(h) F_{ri}(t) + F_{ji}(t)\,. \qquad [3]$$

Note that

$$F_{ji}(t+h) - F_{ji}(t) = P(t \leq H_A < t+h, Y_{t+h} = j | Y_0 = i \in A\,) \qquad [4]$$

is the probability that the transition from $A$ to $j \in B$ occurred within the interval $(t, t+h]$, given the process started in state $i \in A$. Since $F_{jr}(0) = 0$ for all $r \in A$, we may write Eq. [3] as

$$F_{ji}(t+h) - F_{ji}(t) = \sum_{r \in A} \Big(F_{jr}(h) - F_{ri}(0)\Big) F_{ri}(t)\,. \qquad [5]$$

Division by $h$ and taking the limit for $h \to 0$ gives

$$\frac{dF_{ji}(t)}{dt} = \sum_{r \in A} w_{jr} F_{ri}(t)\,, \qquad [6]$$

since

$$\frac{F_{jr}(h) - F_{ri}(0)}{h} \longrightarrow w_{jr}$$

for $h \to 0$; *i.e.*, the rate constants are the time-derivates of the transition functions at time 0 [1]. Summing over all $j \in B$ we obtain with Eq. [4]:

$$P(t \leq H_A < t+h | Y_0 = i \in A) = \sum_{j \in B} \Big(F_{ji}(t+h) - F_{ji}(t)\Big)\,. \qquad [7]$$

The conditional density function of $H_A$, given the process started in $i \in A$, $f_i(t)$, is obtained by dividing Eq. [7] by $h$ and taking the limit for $h \to 0$:

$$f_i(t) = \sum_{j \in B} \frac{dF_{ji}(t)}{dt} = \sum_{j \in B} \sum_{r \in A} w_{jr} F_{ri}(t) \qquad [8]$$

for all $i \in A$, where the second equality follows with Eq. [6].

Let $\boldsymbol{f}_A(t)$ be the *row* vector whose $i^{th}$ component is $f_i(t)$, where $i \in A$. Thus, we may write equations [8] in matrix form:

$$\boldsymbol{f}_A(t) = \boldsymbol{u}_B \mathrm{W}_{BA} F_{AA}(t)\,. \qquad [9]$$

The transition matrix $F(t)$, and thus also $F_{AA}(t)$, is determined by the differential equation:

$$\frac{dF(t)}{dt} = \begin{pmatrix} \mathrm{W}_{AA} & \mathbf{0} \\ \mathrm{W}_{BA} & \mathbf{0} \end{pmatrix} \begin{pmatrix} F_{AA}(t) & F_{AB}(t) \\ F_{BA}(t) & F_{BB}(t) \end{pmatrix} = \begin{pmatrix} \mathrm{W}_{AA} F_{AA}(t) & \mathbf{0} \\ \mathrm{W}_{BA} F_{BA}(t) & \mathbf{0} \end{pmatrix} \; (^3) \qquad [10]$$

and therefore

$$\frac{dF_{AA}(t)}{dt} = \mathrm{W}_{AA} F_{AA}(t)\,. \qquad [11]$$

The solution of the last equation is [1]

$$F_{AA}(t) = e^{t\mathrm{W}_{AA}} = \sum_{k=0}^{\infty} \frac{(t\mathrm{W}_{AA})^k}{k!}\,. \qquad [12]$$

According to the spectral expansion theorem of linear algebra, $\mathrm{W}_{AA}$ may be written as

$$\mathrm{W}_{AA} = \sum_{i \in A} \lambda_i \mathrm{A}_i\,, \qquad [13]$$

where $\lambda_i$ are the eigenvalues of $\mathrm{W}_{AA}$, and $\mathrm{A}_\alpha$ are corresponding $(s \times s)$-matrices, provided $\mathrm{W}_{AA}$ is diagonalizable — *i.e.*, $\mathrm{W}_{AA}$ has $s$ linearly independent eigenvectors. Note that

$$\lambda_i \leq 0 \qquad [14]$$

for all $i \in A$, which is a consequence of Eq. [1] and Gershgorin's "disc theorem" [2].

The matrices $\mathrm{A}_i$ are calculated as follows. Let $B$ be the matrix whose $i^{th}$ column vector, $\boldsymbol{b}_i = (b_{1i}, \dots, b_{ri})^T$, is the eigenvector corresponding to eigenvalue $\lambda_i$ of $\mathrm{W}_{AA}$. Let $C = B^{-1}$ be the inverse of $B$ and $\boldsymbol{c}_i = (c_{i1}, \dots, c_{is})$ its $i^{th}$ row vector. Now,

---

($^3$) Note that for the modified process, $F_{AB}(t) = \mathbf{0}, F_{BB}(t) = \mathbf{0}$ for all $t$.

$$\mathrm{A}_i = \boldsymbol{b}_i \boldsymbol{c}_i \,. \qquad [15]$$

Again, with Eq. [15], it follows that

$$\mathrm{A}_i \mathrm{A}_j = \begin{cases} \mathbf{0} \;\; if \;\; i \neq j \\ \mathrm{A}_i \; if \; i = j \end{cases} . \qquad [16]$$

Relationships [16] imply

$$(\mathrm{W}_{AA})^k = \sum_{i \in A} (\lambda_i)^k \mathrm{A}_i \,. \qquad [17]$$

Insertion of Eq. [17] into [12] gives

$$\begin{aligned} F_{AA}(t) &= \sum_{k=0}^{\infty} \frac{(t\mathrm{W}_{AA})^k}{k!} \\ &= \sum_{\alpha \in A} \left( \sum_{k=0}^{\infty} \frac{(\lambda_i t)^k}{k!} \right) \mathrm{A}_i \qquad [18] \\ &= \sum_{i \in A} e^{\lambda_i t} \mathrm{A}_i \,. \end{aligned}$$

Insertion of this result into Eq. [9] yields

$$\boldsymbol{f}_A(t) = \boldsymbol{u}_B \mathrm{W}_{BA} \left( \sum_{i \in A} e^{\lambda_i t} \mathrm{A}_i \right). \qquad [19]$$

The process begins its sojourn in $A$ only in states that are accessible from $B$. The probability that the process begins in $i \in A$ is the probability that the unmodified process when it leaves $B$ enters $i$. This probability is equal to the probability mass flow per unit time from $B$ into $i \in A$,

$$v_i(t) = \sum_{j \in B} w_{ij} \, p_j(t) \,,$$

divided by the total flow of probability mass per unit time from $B$ into $A$, $\boldsymbol{u}_A \boldsymbol{v}(t)$, where $\boldsymbol{v}(t) = (v_1(t), \ldots, v_s(t))^T$. Thus,

$$\boldsymbol{r}(t) = \frac{\boldsymbol{v}(t)}{\boldsymbol{u}_A \boldsymbol{v}(t)} = \frac{\mathrm{W}_{AB} \boldsymbol{p}_B(t)}{\boldsymbol{u}_A \mathrm{W}_{AB} \boldsymbol{p}_B(t)} \,. \qquad [20]$$

The density function of $H_A$, then, is given by

$$f_A(t) = \boldsymbol{f}_A(t)\boldsymbol{r}(t)\,. \quad [21]$$

Combination of Eq.'s [19], [20], and [21] gives

$$f_A(t) = \boldsymbol{u}_B \mathrm{W}_{BA} \left( \sum_{i \in A} e^{\lambda_i t} \mathrm{A}_i \right) \frac{\mathrm{W}_{AB} \boldsymbol{p}_B(t)}{\boldsymbol{u}_A \mathrm{W}_{AB} \boldsymbol{p}_B(t)}\,. \quad [22]$$

In summary, we may state the following, well-known result [3,4]:

THEOREM 1. For a time-homogeneous Markov process, the probability density function of $H_A$ — the dwell time of the process in any subset of states $A$ — is given by the linear combination of decaying exponentials

$$f_A(t) = \sum_{i \in A} c_i(t) e^{\lambda_i t}\,, \quad [23]$$

for all $i \in A$, where the $\lambda_i's \leq 0$ are the eigenvalues of the generator submatrix $\mathrm{W}_{AA}$, which must be diagonalizable, and coefficients

$$c_i(t) = \boldsymbol{u}_B \mathrm{W}_{BA} \mathrm{A}_i \frac{\mathrm{W}_{AB} \boldsymbol{p}_B(t)}{\boldsymbol{u}_A \mathrm{W}_{AB} \boldsymbol{p}_B(t)}\,. \quad [24]$$

Table S3 (see below) lists exponential coefficients for different graphs.

COMMENT 1. The submatrix $\mathrm{W}_{AA}$ only contains rate constants for transitions between $A$ states and from $A$ states into $B$ states. The eigenvalues of $\mathrm{W}_{AA}$, $\lambda_i$, are therefore functions of these rate constants, and not others.

COMMENT 2. For steady state (stationary) processes, the coefficients $c_i$ become time-independent, for $\boldsymbol{p}_B(t)$ in Eq. [20] is replaced by $\boldsymbol{p}_B$.

COMMENT 3. For the sake of calculating the dwell time distribution of the sojourn in $A$, $B$-states may be replaced by a single state with rate constant $v_i(t)$ for the transition out of $B$ into state $i \in A$. In this instance, the probabilities $p_{j \in B}(t)$ are absorbed into the rate constant $v_i(t)$.

CORROLARY 1. The dwell time of a Markov process in single microstates is exponentially distributed.

*Proof*: Let $A = \{1\}$. Then, $\boldsymbol{u}_B \mathrm{W}_{BA} = -w_{11}$, $\mathrm{W}_{AA} = w_{11}$, $\mathrm{A}_1 = 1$, $\lambda_1 = w_{11}$, and $\boldsymbol{u}_A = 1$. With Theorem 1 (Eq.'s [23], [24]) it follows that

$$f_A(t) = -w_{11} e^{w_{11} t},$$

the density of the exponential distribution [5]. ■

In the following we limit our discussion to processes on graphs with three nodes $S = \{1,2,3\}$, $A = \{1,2\}$ and $B = \{3\}$. In general, the generator of the process is

$$\mathrm{W} = \begin{pmatrix} w_{11} & w_{12} & w_{13} \\ w_{21} & w_{22} & w_{23} \\ w_{31} & w_{32} & w_{33} \end{pmatrix}.$$

According to Eq. [23], the density function of $H_A$ is given by

$$f_A(t) = c_1 e^{\lambda_1} + c_2 e^{\lambda_2} . \qquad [25]$$

The exponential parameters, $\lambda_1$ and $\lambda_2$, are the eigenvalues of

$$\mathrm{W}_{AA} = \begin{pmatrix} w_{11} & w_{12} \\ w_{21} & w_{22} \end{pmatrix}.$$

In the general case,

$w_{11} = -w_{21} - w_{31}$ and $w_{22} = -w_{12} - w_{32}$ (*cf.* Eq. [1]).

The eigenvalues of $\mathrm{W}_{AA}$ are the roots of its characteristic polynomial. In the general case, the eigenvalues are

$$\begin{aligned}\lambda_1 = &-\frac{1}{2}(w_{12} + w_{21} + w_{31} + w_{32}) \\ &-\frac{1}{2}\sqrt{(w_{12} + w_{21} + w_{31} + w_{32})^2 - 4(w_{12}w_{31} + w_{21}w_{32} + w_{31}w_{32})}\end{aligned} \qquad [26]$$

and

$$\begin{aligned}\lambda_2 = &-\frac{1}{2}(w_{12} + w_{21} + w_{31} + w_{32}) \\ &+\frac{1}{2}\sqrt{(w_{12} + w_{21} + w_{31} + w_{32})^2 - 4(w_{12}w_{31} + w_{21}w_{32} + w_{31}w_{32})}\end{aligned} \qquad [27]$$

The coefficients $c_1$ and $c_2$ are cumbersome expressions in general. However, the sum of the two is surprisingly simple:

$$c_1 + c_2 = \frac{w_{13}w_{31} + w_{23}w_{32}}{w_{13} + w_{23}}. \qquad [28]$$

LEMMA 1. For the Sequence and Cycle 1 (*cf.* Fig. 5A), the forward rate constants $w_{32}$ and $w_{21}$, are bounded by $\lambda_1$ and $\lambda_2$:

$$-\lambda_2 < w_{32}, w_{21} < -\lambda_1 \,.$$

*Proof*: To simplify notation, let $a = w_{12}$ , $b = w_{21}$, and $x = w_{32}$. Since $a, b > 0$,

$$b < a + b \,,$$

which implies the following inequalities:

$$4bx < 4(a+b)x$$

$$0 < 4(a+b)x - 4bx$$

$$-2(a+b)x < 2(a+b)x - 4bx$$

$$(a+b)^2 - 2(a+b)x + x^2 < (a+b)^2 + 2(a+b)x - 4bx + x^2$$

$$(a+b-x)^2 < (a+b+x)^2 - 4bx$$

$$a+b-x < \sqrt{(a+b+x)^2 - 4bx}$$

$$a+b+x < \sqrt{(a+b+x)^2 - 4bx} + 2x$$

$$\frac{1}{2}(a+b+x) - \frac{1}{2}\sqrt{(a+b+x)^2 - 4bx} = -\lambda_2 < x \,,$$

where the equality follows from Eq. [27] and $w_{31} = 0$.

Likewise, with $(a+b-x)^2 = (x-a-b)^2$ it follows from the fifth line above that

$$x - (a+b) < \sqrt{(a+b+x)^2 - 4bx}$$

$$2x - (a+b) < x + \sqrt{(a+b+x)^2 - 4bx}$$

$$x < \frac{1}{2}(a+b+x) + \frac{1}{2}\sqrt{(a+b+x)^2 - 4bx} = -\lambda_1 \,,$$

where the equality is due to Eq. [26] and $w_{31} = 0$. Thus,

$$-\lambda_2 < w_{32} < -\lambda_1.$$

Multiplication of the last inequality with $w_{21} > 0$ yields

$$-\lambda_2 w_{21} < w_{21} w_{32} < -\lambda_1 w_{21}$$

and with $\lambda_1\lambda_2 = w_{21}w_{32}$ it follows that

$$-\lambda_2 < w_{21} < -\lambda_1 \, . \blacksquare$$

THEOREM 2. For trees (Fork and Sequence, *cf.* Fig. 5), $c_1$, $c_2 > 0$ in Eq. [27].

*Proof*: The coefficients are given by Eq. [24]. In case of the Fork,

$$\boldsymbol{r}(t) = \frac{1}{w_{13} + w_{23}} \begin{pmatrix} w_{13} \\ w_{23} \end{pmatrix},$$

$$A_1 = \begin{pmatrix} 1 & 0 \\ 0 & 0 \end{pmatrix},$$

$$A_2 = \begin{pmatrix} 0 & 0 \\ 0 & 1 \end{pmatrix},$$

$\boldsymbol{u}_B = 1$, $\mathrm{W}_{BA} = (w_{31}, w_{32})$, $\lambda_1 = -w_{31}$ and $\lambda_2 = -w_{32}$ (*cf.* Eq.'s, [26] and [27])

$$c_1 = \boldsymbol{u}_B \mathrm{W}_{BA} \mathrm{A}_1 \boldsymbol{r}(t) = -r_1(t)\lambda_1 > 0 \, ,$$

$$c_2 = \boldsymbol{u}_B \mathrm{W}_{BA} \mathrm{A}_2 \boldsymbol{r}(t) = -r_2(t)\lambda_2 > 0 \, .$$

The inequality sign follows with [14]. (We kept the time dependence for $\boldsymbol{r}$ and its components, $r_i$, because if State 3 is a compound state, the rate constants become functions of time, unless the process is in steady state, *cf.* Comment 3.)

In case of the Sequence,

$$\boldsymbol{r}(t) = \begin{pmatrix} 0 \\ 1 \end{pmatrix},$$

$$A_1 = -\frac{1}{\lambda_2 - \lambda_1} \begin{pmatrix} \lambda_2 + w_{21} & w_{12} \\ w_{21} & \lambda_1 + w_{21} \end{pmatrix},$$

$$A_2 = \frac{1}{\lambda_2 - \lambda_1} \begin{pmatrix} -\lambda_1 - w_{21} & w_{12} \\ w_{21} & \lambda_2 + w_{21} \end{pmatrix},$$

$\boldsymbol{u}_B = 1$ and $\mathrm{W}_{BA} = (0, w_{32})$. Thus, with $\lambda_1\lambda_1 = w_{32}w_{21}$, it follows that

$$c_1 = \boldsymbol{u}_B \mathrm{W}_{BA} \mathrm{A}_1 \boldsymbol{r} = -\frac{w_{32}(\lambda_1 + w_{21})}{\lambda_2 - \lambda_1} = -\frac{w_{32} + \lambda_2}{\lambda_2 - \lambda_1}\lambda_1$$

and

$$c_2 = \boldsymbol{u}_B \mathrm{W}_{BA} \mathrm{A}_2 \boldsymbol{r} = \frac{w_{32}(\lambda_2 + w_{21})}{\lambda_2 - \lambda_1} = \frac{w_{32} + \lambda_1}{\lambda_2 - \lambda_1} \lambda_2 \,.$$

With Lemma 1 it follows that $w_{32} + \lambda_2 > 0$ and $w_{32} + \lambda_1 < 0$. Hence, with [14] follows that $c_1, c_2 > 0$. This proves the assertion. ■

THEOREM 3. For Cycle 1 (*cf.* Fig. 5), the coefficients of the dwell time density function are either all positive or mixed (positive and negative).

*Proof*: The coefficients of the density function for the dwell time in the compound state are

$$c_1 = -\frac{\lambda_2(w_{13} + w_{23}) + w_{23}w_{32}}{(w_{13} + w_{23})(\lambda_2 - \lambda_1)} \lambda_1 \,, \qquad [29]$$

$$c_2 = \frac{\lambda_1(w_{13} + w_{23}) + w_{23}w_{32}}{(w_{13} + w_{23})(\lambda_2 - \lambda_1)} \lambda_2 \,. \qquad [30]$$

Suppose $c_1, c_2 > 0$, then

$$\lambda_2(w_{13} + w_{23}) + w_{23}w_{32} > 0$$

and

$$\lambda_1(w_{13} + w_{23}) + w_{23}w_{32} < 0 \,,$$

*i.e.*,

$$-\lambda_2 < \frac{w_{23}w_{32}}{w_{13} + w_{23}} < -\lambda_1 \,.$$

These inequalities may apply, if $w_{23}/(w_{13} + w_{23})$, the probability of entering the compound state *via* State 2 rather than 1, is sufficiently close to 1 (*cf.* Lemma 1). However, for sufficiently small probabilities of beginning the sojourn in the compound state in State 2 rather than 1,

$$w_{23}w_{32}/(w_{13} + w_{23}) < -\lambda_2.$$

In this case, $c_1 < 0$ (especially when $w_{23} = 0$, for in this case the sojourn in the compound state always begins in state 1 and not 2). ■

THEOREM 4. Only for Cycle 2 is

$$c_1 + c_2 = 0.$$

*Proof*: Cycle 2 fulfills $c_1 + c_2 = 0$ (*cf.* Table S3). Now let $c_1 + c_2 = 0$. By integrating Eq. [25], putting $t = 0$, and observing that $F_A(0) = 0$, we obtain $c_1 = \lambda_1\lambda_2/(\lambda_1 - \lambda_2) < 0$, the result obtained for Cycle 2 (*cf.* Table S3). Alternatively, with Eq. [28] it is seen that $c_1 + c_2 = 0$ if and only if $w_{13}w_{31} + w_{23}w_{32} = 0$. Only two ergodic graphs fulfill this requirement: the graph with $w_{13}, w_{32} = 0$ and the graph with $w_{31}, w_{23} = 0$. Both are isomorphic to Cycle 2. ■

THEOREM 5. For the Sequence, Cycle 1 and Cycle 2 (*cf.* Fig. 5), if

$$w_{21}w_{32} \to 0, \qquad \text{then} \qquad c_2 \to 0.$$

*Proof*: For all three graphs, $w_{31} = 0$. Insertion of $w_{31} = 0$ into Eq. [27] shows that $\lambda_2 \to 0$ if $w_{21}w_{32} \to 0$. The claim now follows with Eq. [30], which also applies to Cycle 2 and the Sequence because both are obtained from Cycle 1 by pruning (i.e., by setting specific rate constants to zero). ■

#### Autocorrelation Functions

The autocorrelation function for the signal generated by a stationary stochastic process gives the signal correlation for pairs of timepoints, $t_1 \le t_2$, as a function of $\tau = t_2 - t_1$; the difference, $\tau$, is called the lag-time. Loosely speaking, the autocorrelation function indicates for how long, on average, the presence holds sway over the future, or how fast the process "forgets" its past. For a random telegraph process with forward rate $w_{21}$ and reverse rate $w_{12}$ that generates binary sample paths alternating between 0 and 1, the autocorrelation function is given by

$$acf(\tau) = e^{-\kappa\tau}, \qquad [31]$$

where $\kappa = w_{12} + w_{21}$ [5]. The average length of 0-signal (OFF) and 1-signal (ON) periods is given by $w_{21}^{-1}$ and $w_{12}^{-1}$, respectively.

Although incorrect, the random telegraph model may serve the goal of elucidating the relationship between period length and autocorrelation. Eq. [31] shows that the autocorrelation function is highly sensitive to the average length of both ON and OFF periods: a shortening of periods shifts the function toward shorter lag-times, extension of lag-times toward longer lag-times. If ON periods are significantly shorter than OFF periods, as observed, then the $acf$ is dominated by burst length, *i.e.*, by the magnitude of $w_{12}$, and not burst frequency, $w_{21}$. Thus, small changes in $w_{12}$ may have a larger effect on $acf$ than large changes in $w_{21}$, which may explain why the autocorrelation functions of wild type and activator mutant were nearly identical (*cf*. Fig. S1), despite the marked decrease in burst frequency in the activator mutant (*cf*. Fig. 1). The same argument applies to the *chd1*Δ and *isw2*Δ mutant. With the average lengths of ON and OFF periods provided in Tables S1 and S2, the random telegraph model predicts closely similar autocorrelation functions for all strains analyzed.

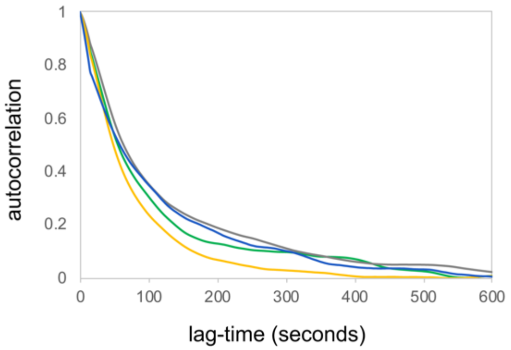


**Fig. S1.** ([4])

A comparison of the autocorrelation function calculated from fluorescence sample paths and binary sample paths, which were obtained from the former by segmentation into ON and OFF periods and setting the signal during ON and OFF periods to 1 and 0, respectively, demonstrates that both are closely similar for all strains analyzed (Fig. S2), indicating that autocorrelation functions from fluorescence sample paths were largely determined by the length of ON and OFF periods.

[4] **Fig. S1.** Autocorrelation functions from fluorescence sample paths for "wild type" (*pho80*Δ*,* gray), activator mutant (*pho4*Δ[85-99], blue), *chd1*Δ mutant (green), and *isw2*Δ mutant (yellow).

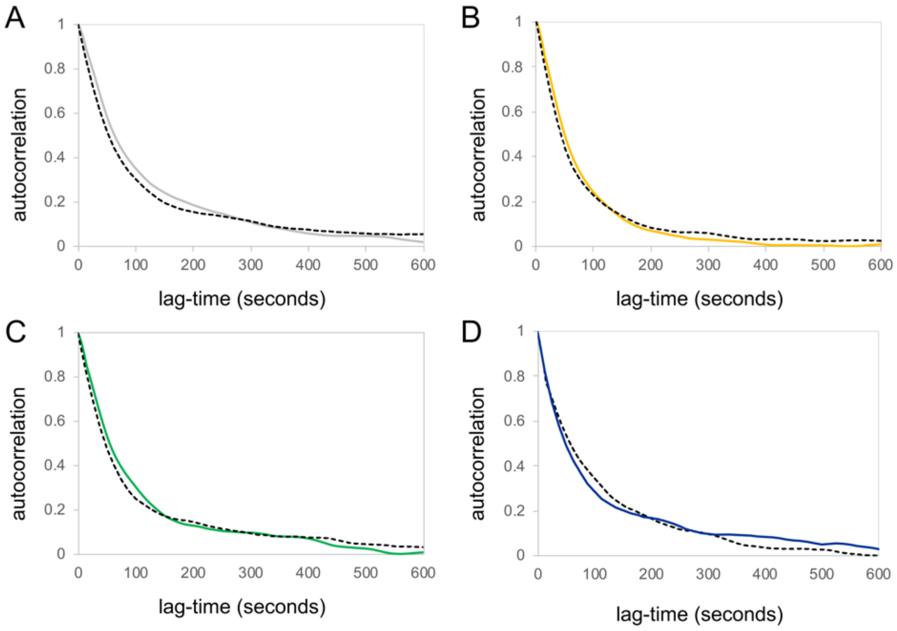


**Fig. S2.** ([5])

| | $\lambda_1\ (s^{-1})$ | $\lambda_2\ (s^{-1})$ | $c_1\ (s^{-1})$ | $c_2\ (s^{-1})$ | $mean\ (s)$ |
|---|---|---|---|---|---|
| WT | −0.049 | −0.0128 | −0.0173 | 0.0173 | 98 |
| pho4Δ [75-90] | −0.101 | −0.014 | −0.0162 | 0.0162 | 81 |
| isw2Δ | −0.050 | −0.015 | −0.0219 | 0.0219 | 85 |
| chd1Δ | −0.066 | −0.013 | −0.0168 | 0.0168 | 89 |

**Table S1.** ([6])

[5] **Fig. S2.** Comparison of autocorrelation functions from fluorescence and binary sample paths. The latter were obtained from the former by segmentation into ON and OFF periods and setting the signal for ON and OFF periods to 1 and 0, respectively. The graphs of autocorrelation functions from fluorescence sample paths are shown as continuous lines, the corresponding autocorrelation functions from binary sample paths are shown as dashed black lines. (A) Wild type. (B) The *isw2*Δ mutant (yellow). (C) The *chd1*Δ mutant. (D) The *pho4*Δ[85-99] mutant.

[6] **Table S1.** Probability density functions for observed ON periods. Densities were obtained by taking the derivative of the best-fit of $F(t) = 1 + ae^{\lambda_1 t} - (1 + a)e^{\lambda_2 t}$ to the observed distribution function. Densities have the form $f(t) = c_1 e^{\lambda_1 t} + c_1 e^{\lambda_2 t}$, where $c_1 = a\lambda_1$ and $c_2 = -(1 + a)\lambda_2$.

| $\times 10^{-3}$ | $\lambda_1 (s^{-1})$ | $\lambda_2\ (s^{-1})$ | $c_1\ (s^{-1})$ | $c_2\ (s^{-1})$ | $mean\ (s)$ |
|---|---|---|---|---|---|
| WT | −3.6 | −1.15 | 1.825 | 0.566 | 570 |
| pho4Δ [75-90] | −0.588 | − | 0.588 | − | 1665 |
| isw2Δ | −3.65 | −1.04 | 1.286 | 0.672 | 720 |
| chd1Δ | −2.63 | −0.588 | 1.926 | 0.158 | 734 |

**Table S2.** ([7])

| $f(t) = c_1 e^{\lambda_1 t} + c_2 e^{\lambda_2 t}$ | $c_1$ | $c_2$ |
|---|---|---|
| Fork | $-\dfrac{w_{13}\lambda_1}{w_{13}+w_{23}}$ | $-\dfrac{w_{23}\lambda_2}{w_{13}+w_{23}}$ |
| Sequence | $-\dfrac{w_{32}+\lambda_2}{\lambda_2-\lambda_1}\lambda_1$ | $\dfrac{w_{32}+\lambda_1}{\lambda_2-\lambda_1}\lambda_2$ |
| Cycle 1 | $-\dfrac{\lambda_2(w_{13}+w_{23})+w_{23}w_{32}}{(w_{13}+w_{23})(\lambda_2-\lambda_1)}\lambda_1$ | $\dfrac{\lambda_1(w_{13}+w_{23})+w_{23}w_{32}}{(w_{13}+w_{23})(\lambda_2-\lambda_1)}\lambda_2$ |
| Cycle 2 | $-\dfrac{\lambda_2\lambda_1}{\lambda_2-\lambda_1}$ | $\dfrac{\lambda_1\lambda_2}{\lambda_2-\lambda_1}$ |

**Table S3.** ([8])

### Citations


1. Cinlar, E. *Introduction to stochastic processes*. (Dover Publications, 2013).

2. Mirzaev, I. & Gunawardena, J. Laplacian Dynamics on General Graphs. *Bull. Math. Biol.* **75**, 2118–2149 (2013).

3. D . Colquhoun and A . G . Hawkes Source. On the Stochastic Properties of Single Ion Channels. *Proc. R. Soc. London . Ser. B, Biol. Sci.* **211**, 205–235 (1981).

4. Tu, Y. The nonequilibrium mechanism for ultrasensitivity in a biological switch: Sensing by Maxwell's demons. *Proc. Natl. Acad. Sci. U. S. A.* **105**, 11737–11741 (2008).


[7] **Table S2.** Probability density functions for observed OFF periods. Densities were obtained by taking the derivative of the best-fit of $F(t) = 1 + ae^{\lambda_1 t} - (1+a)e^{\lambda_2 t}$ to the observed distribution function. Densities have the form $f(t) = c_1 e^{\lambda_1 t} + c_1 e^{\lambda_2 t}$.

[8] **Table S3.** Analytic expressions of the probability density functions for the dwell time of a Markov process in the compound ON/OFF state for different transition topologies. The compound state encompasses two microstates.

5. Grimmet, G. & Stirzaker, D. *Probability and random processes*. (Oxford university press, 2001).
